# Read coverage as an indicator of misassembly in a short-read based genome assembly

**DOI:** 10.1101/790337

**Authors:** Peipei Wang, Fanrui Meng, Bethany M. Moore, Shin-Han Shiu

## Abstract

Availability of genome sequences has led to significant advance in biology. With few exceptions, the great majority of existing genome assemblies are derived from short read sequencing technologies with highly uneven read coverages indicative of sequencing and assembly issues. In tomato, 0.6% (5.1 Mb) and 9.7% (79.6 Mb) of short-read based assembly had significantly higher and lower coverage compared to background, respectively. We established machine learning models capable of predicting genomic regions with variable coverages and found that high coverage regions tend to have lower simple sequence repeat but higher tandem gene densities compared to background regions. To determine if the high coverage regions were misassembled, we examined a recently available long-read based assembly and found that 27.8% (1.41 Mb) of high coverage regions were potentially mis-assembled of duplicate sequences, compared to 1.4% in background regions. In addition, using a machine learning model that can distinguish correctly and incorrectly assembled high coverage regions, we found that misassembled, high coverage regions tend to be flanked by simple sequence repeats, pseudogenes, and transposon elements. Our study provides insights on the causes of variable coverage regions and a quantitative assessment of factors contributing to misassembly when using short reads.

## INTRODUCTION

The number of whole genome sequences has increased dramatically in the last decades due to the development of new generations of sequencing technologies and reduced cost. The “first” generation was Sanger sequencing technology (1), based on which a decade was taken to deliver a draft genome of human (2). The second-generation technology—i.e., “Next generation sequencing” where thousands to millions of DNA molecules are enabled to be sequenced simultaneously dramatically shortens the time required to obtain high genome coverage (1). However, due to the short length of these reads (36bp~400bp), there are many challenges for assembling genome based on short reads, including the difficulty in sequencing repetitive sequences (3), low read coverages in GC-poor or GC-rich regions (4), genome sequencing bias introduced by PCR amplification during library construction (5), and polyploidy in some species including most flowering plants (6). The advent of third generation sequencing, e.g., Pacific Biosciences (PacBio) single molecule real time sequencing (7) and Oxford Nanopore sequencing (8), has led to another revolution in genome sequencing, where long reads up to 100 Kb can be sequenced in a single run without PCR amplification or chemical labeling of the sample. Although the much higher error rates remain an issue (9), the third generation sequencing still has merit for applicants more tolerant to error rates, like structural variant calling (10) and, combined with short-read sequencing, is overtaking projects that focus on short reads only.

Although the number of genome sequences which take the advantages of both second and third generation sequencing are increasing (11–13), the majority of genome assemblies available from the National Center of Biotechnology Information (NCBI) were generated predominantly using short reads from the second-generation technology. Before the third-generation sequencing is more widely applied to improve these genome assemblies, it remains important to assess the quality of existing short-read based assemblies. Several methods have been developed for this purpose, like Scaffold N50, MaGuS (14), LTR Assembly Index (15), SQUAT (16). In addition to these methods, another strategy is to assess how well an assembly is covered by the reads used for building the assembly.

For an ideal genome assembly, the sequencing reads would be uniformly distributed across the genome. However, in the real world, when sequencing reads are mapped back to the genome, the read coverage varies across genome due to multiple reasons. First, regions with extremely high or low GC content may not be sequenced equally compared with other GC-balanced regions, leading to low or even no coverage of reads (17). Second, repetitive sequences are abundant in species with larger genomes, and have always been a major challenge for genome assemblies (3). Repeats longer than read length would lead to gaps in the genome assembly due to uncertainty in assembly of these regions. This would break down the genome into pieces, leading to the loss of linkage information among genetic markers. Third, repeats may also be led to misassembly where two unlinked regions were joined together and resulted in higher than usual read coverages. In the case of repetitive sequences containing genes, such as tandemly duplicated genes and retrogenes, such misassembly would reduce the gene copy number estimation. These missing genes not only make it challenging to account for all the genes in a genome but also create problems for functional genomic studies by impacting gene expression level estimates or loss-of-function studies. For example, the annotated SEC10a gene from *Arabidopsis thaliana* and its recent tandem duplicate copy SEC10b, were assembled together and annotated as a single gene, which explains why homozygous T-DNA insertion mutant of either copy has no phenotypic change compared to wild-type (18).

Here, we use tomato (*Solanum lycopersicum*) as a model to assess the extent to which its assembly has variable coverage. Tomato is chosen because assemblies built with short reads, as well as PacBio long reads were both available. In addition, it is an important crop and a major model species for studying specialized metabolism, particularly considering specialized metabolism genes tend to be duplicated tandemly (19,20) and may tend to be misassembled. The idea of searching for regions with significantly high or low read coverages has been extensively applied in estimating copy number variation (CNV) among species or populations (21–23). Using CNV detection tools, we are interested in identifying genome regions with significantly high- and low-coverage of sequencing reads compared to the genome average (i.e. background). Most importantly, based on information on regions with variable coverage, our primary goals are to explore underlying factors influencing the read coverages and, most importantly, assembly quality through comparison between short and long-read assemblies.

## MATERIALS AND METHODS

### Genome assembly and sequencing reads

The genome sequence assembly SL2.50 of tomato cultivar ‘Heinz 1706’ was downloaded from NCBI (https://www.ncbi.nlm.nih.gov/). The genome was assembled mainly using 454 reads, Sanger sequencing reads of two sets of Bacterial Artificial Chromosome (BAC) clone pools, and BAC and Fosmid clone end sequences (24), and was referred to as the Short-read assembly. Additional SOLiD and Illumina reads were used for the base error correction. Among the scaffolds, 91 of 3223 were anchored to 12 chromosomes (25). To evaluate the extent of mis-assembly, tomato assembly SL4.0, which was assembled using 80X PacBio sequences (referred to as the Long-read assembly), was also downloaded from Solanaceae Genomics Network (SGN, https://www.solgenomics.net).

There are two batches of Illumina genomic sequencing reads available in tomato. The first batch of reads (referred to as dataset1), used for base error correction in genome assembly with a ~28-fold coverage of the genome (24), was sequenced with an Illumina Genome Analyzer IIx (GAIIx) sequencer, and obtained from the SGN in the form of BAM file (version SL2.9). The other read batch (dataset2) was sequenced with an Illumina HiSeq 2000 sequencer, with a ~46-fold coverage of the genome (SRP010718). Reads of these two datasets were remapped to the Short-read assembly and Long-read assembly using Burrows-Wheeler Aligner (BWA-MEM) (26) with default parameters. To eliminate the impact of bias in PCR amplification on read coverage calling, duplicate reads (identical reads with same mapped location) were marked and removed using Picard (http://broadinstitute.github.io/picard). Due to concern of data quality (see **Results**), dataset1 was not analyzed further.

### Estimation of read depth and detection of variable coverage regions

Regions with high/low read coverage were identified using CNVnator (21) by determining Read Depth (RD) for an optimally sized bin of the genome assembly as the number of mapped reads with ≥50% of read lengths overlapping with the bin boundaries (21). The optimal bin size was the bin size leading to a ratio of RD average to RD standard deviation of ~4-5 as suggested (21). For dataset2, bin sizes from 50bp to 300bp were evaluated, and 100bp was chosen with a ratio = 5.18 (**Supplementary Table S1**). With bin size of 100bp, 20,071 and 1,385 regions were identified in Short-read assembly as low-coverage (LC) and high-coverage (HC) regions by CNVnator, respectively. The remaining 20,743 regions were treated as background (BG). The RD values from this CNVnator run for further analysis is referred to as “analysis RD” values.

To assess the sensitivity and accuracy of HC/LC region detection, reads were resampled with three strategies to generate simulated RD values (for details, see **Supplemental Text**) that were used to run CNVnator. For each strategy, the resultant RD for each 100bp bin was compared to the simulated RD values (ground truth). In the first strategy, reads were resampled for HC, BG and LC regions, based on idealized RD values (input RDs for HC, BG and LC regions were assigned as 2, 1 and 0, respectively, no decimal point values). In the second strategy, reads were resampled for HC, BG and LC regions based on the rounded analysis RD values (e.g., for regions with analysis RD values of 0.88 and 2.31, 1x and 2x reads were resampled for these two regions, respectively). In the third strategy, reads were resampled for HC, BG and LC regions based on the analysis RD values. For all three strategies, we observed very high correlation between simulated, ground truth RD and RD values resulted from simulated reads, indicating that detection of HC/LC/BG regions using CNVnator is robust.

### Genome and functional annotations, definitions of genome features, and gene set enrichment analysis

Tomato gene annotation version SL2.50 was downloaded from NCBI (https://www.ncbi.nlm.nih.gov/). Aside from gene annotations, we defined or obtained additional genome features including pseudogenes, transposable elements (TE), simple sequence repeat (SSR; stretch of DNA, 2 ~ 64bp, repeated >1 time and the repetitions are immediately adjacent to each other), and tandemly duplicated genes. Pseudogenes were defined as genomic regions with significant similarity to protein-coding genes had premature stops/frameshifts and/or were truncated as described in (27). Transposable element (TE) annotation was based on SGN ITAG2.4 release. SSRs were detected using Tandem Repeats Finder with recommended parameters with Match = 2, Mismatch = 7, Delta = 7, PM = 80, PI = 10. Minscore = 50, Maxperiod = 500 (28). Tandemly duplicated genes were identified using MCScanX-transposed (29), as described previously (27), where paralogs are directly adjacent to each other, or separated by ≤10 nonhomologous genes.

Three types of functional annotation data were used including Gene Ontology (GO) terms, Metabolic pathway annotation, and Pfam domain annotation. GO terms were inferred using blast2go (30), where protein sequences were searched against NCBI nr protein database using BLASTP (31) with an E-value cut-off of 1e-5. Tomato metabolic pathway annotation V3.0 was downloaded from Plant Metabolic Network (https://pmn.plantcyc.org/). Genes in specialized metabolic pathways were annotated as specialized metabolism (SM) genes. Pfam domains in tomato annotated protein sequences were identified by searching against Pfam Hidden Markov Models (https://pfam.xfam.org, v.29.0) using HMMER3 (http://hmmer.org) with the trusted cutoff.

Gene set enrichment analysis was performed using Fisher’s exact test and Likelihood Ratio test, p-values were adjusted for multiple testing (32). Log likelihood ratio was calculated as: 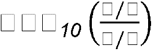, where *a*, *b*, *c* and *d* are the numbers in a 2×2 contingency table as in Fisher’s exact test. For example, for testing whether genes with the GO term *G* tend to be in HC region: *a* -the number of genes with GO term *G* in HC regions, *b* -the number of genes that don’t have GO term *G* but are in HC regions, *c* -the number of genes with GO term *G* but not in HC regions, and *d* -the number of genes that don’t have GO term *G* and are not in HC regions.

### Multi-class machine learning models for predicting whether a genomic region has high, low, or background coverage

To classify a genomic region into one of the following three classes: HC, LC, and BG, three-class models were established using Random Forest (33), implemented in the python package Scikit-learn (34). ~7,000 properties of genomic sequences within HC, LC and BG regions, and in their corresponding flanking regions (in bins of 0.5, 1, 2, 4, 8, 16, and 32 Kb, including both upstream and downstream) were used as features for building machine learning models. There were three types of features. The first was GC content. The second includes densities of: (1) all genes, (2) tandemly duplicated genes, (3) non-tandem genes, (4) pseudogenes, (5) transposable elements, (6) all SSRs (without consider each, specific SSR sequence), (7) specific SSR (2-64bp repeats, e.g. 2bp repeat: ATATATATATAT), and (8) specific *k*-mer (1-6bp, e.g. 5-mer, GGCGG). Density was calculated as the proportion of each HC/LC/BG region occupied by the feature in question. The last type was presence/absence of genes with a particular functional annotation in a given region. These functional annotations include: GO term, Pfam domain, and metabolic pathway annotations. In addition to presence/absence, numbers of annotated entries were also used as features to build models, which didn’t differ significantly in performance from models built with presence/absence features and were not discussed further. Kruskal-Wallis H test was done to determine if there are statistically significant differences of each density among HC/LC/BG regions using SciPy (35). Feature selection was conducted using the RandomForestClassifier function in Scikit-learn (34), and potentially informative features were selected based on their importance determined by the entropy criterion which measures the quality of tree split according to the information gain when each feature was used. For each class, 10% of the regions were held out from the model training/validation process to serve as independent test data. The rest 90% were used as training/validation data for model training.

For model training, we used equal numbers of instances from each class (HC, LC, or BG) to create balanced datasets that facilitate interpretation of model performance. Because HC regions were in the minority (1,156), LC and BG regions were randomly sampled till they were the same numbers as HC regions. In total, 100 random balanced datasets were generated. Using each of the first 10 balanced dataset, the grid search approach as implemented in the GridSearchCV function in Scikit-Learn was used to determine the best combination of parameters. In this approach, each balanced dataset was split into training (90%) and validation (10%) subsets following the 10-fold cross-validation scheme. The best set of parameters for Random Forest (*max_depth, max_feature*, and *n_estimators*) were identified according to the mean of F1-macro (average F1 score for three classes) across the 10 GridSearchCV runs. Then each of the 100 balanced datasets was used to establish a three class (HC/LC/BG) prediction model using the best parameter set with 10-fold cross validation to assess the robustness of classification results. The average true positive rate and the average F1 score across 100 runs were used to evaluate model performance.

### Identification of potentially mis-assembled regions

The Short-read assembly (query) was aligned to the Long-read assembly (subject) with the NUCmer algorithms using default settings (--mumreference --delta --breaklen 200 – mincluster 65 –diagfactor 0.12 –maxgap 90 –minmatch 20), as implemented in MUMmer 4.0.0beta2 (36). Chromosome coordinates and sequence similarity of aligned regions were produced by mummerplot utility. Only aligned regions with identities ≥ 95% were used for the downstream analysis which may lead to false negatives.

Before identifying if a Short-read assembly region is mis-assembled, we first ask, in the MUMmer alignment, if a query region of the Short-read assembly was duplicated in the Long-read assembly or not. A query region was classified as having duplicated subjects if it had ≥ two aligned subject regions in the Long-read assembly, regardless of these regions are on the same chromosome or not. Otherwise, it is regarded as non-duplicated. A query region without duplicated subject was defined as mis-assembled if the subject region it aligned to was in a different location on the same or on different chromosome compared to the location of query region. A query region with duplicated subject regions was defined as mis-assembled if it was: 1) ≥ 100bp, and 2) the lengths of overlaps between duplicated subject regions < 50% of the query, and 3) the subject regions aligned to only one Short-read assembly region. We considered mis-assembled regions that had one copy in Short-read assembly while ≥ 2 in Long-read assembly. Thus, mis-assembled regions with ≥ 2 copies in Short-read assembly and even more copies in Long-read assembly were not analyzed because they were minor cases and the challenges in defining additional unifying categories for these cases.

## RESULTS & DISCUSSION

### Abundance of tomato genomic regions with higher and lower than average coverage

Two datasets were used to determine how well different genomic regions were covered and to define variable coverage regions: genomic regions with significantly higher coverage (HC) or lower coverage (LC) than average (referred to as background, BG). The first, dataset1, was generated with Illumina Genome Analyzer IIx (GAIIx) sequencer with 90-bp paired-end and 54-bp mate pair reads (~28x coverage) used in the original genome assembly (24), and the second, dataset2, was generated with Illumina HiSeq 2000 sequencer with 101-bp paired-end reads (~46-fold coverage). To assess the qualities of these two datasets and to see if both datasets should be analyzed, the tomato genome assembly was split into 100bp bins and, for each bin, the read depth (RD) was determined using either dataset1 or 2 (see **Methods**). After correcting the RD values for GC content bias (21), the median RDs for dataset1 and dataset2 are 1.04 and 1.05, respectively (an example region on chromosome 1, Fig. 1A). The RD values from these two datasets are significantly correlated (Spearman’s ρ = 0.40, *p <*2.2e-16), and consistently revealed bins with substantial deviations from the median value in both directions indicating the presence of HC and LC regions (e.g. grey and black arrows in Fig. 1A). However, RDs of dataset1 was significantly more variable across genome (variance = 0.25 using 0~99 percentile values) compared to those of dataset2 (variance = 0.15, F-test, *p* < 2.2e-16, Fig. 1A). This is not simply due to the higher genome coverage of dataset2 (~46.3x) compared to dataset1 (~28.6x), because a subset of randomly sampled reads from dataset2 to ~30x genome coverage has much more similar RD estimates as dataset2 (Spearman’s ρ = 0.87, *p <*2.2e-16, Fig. 1A).

**Figure 1.**
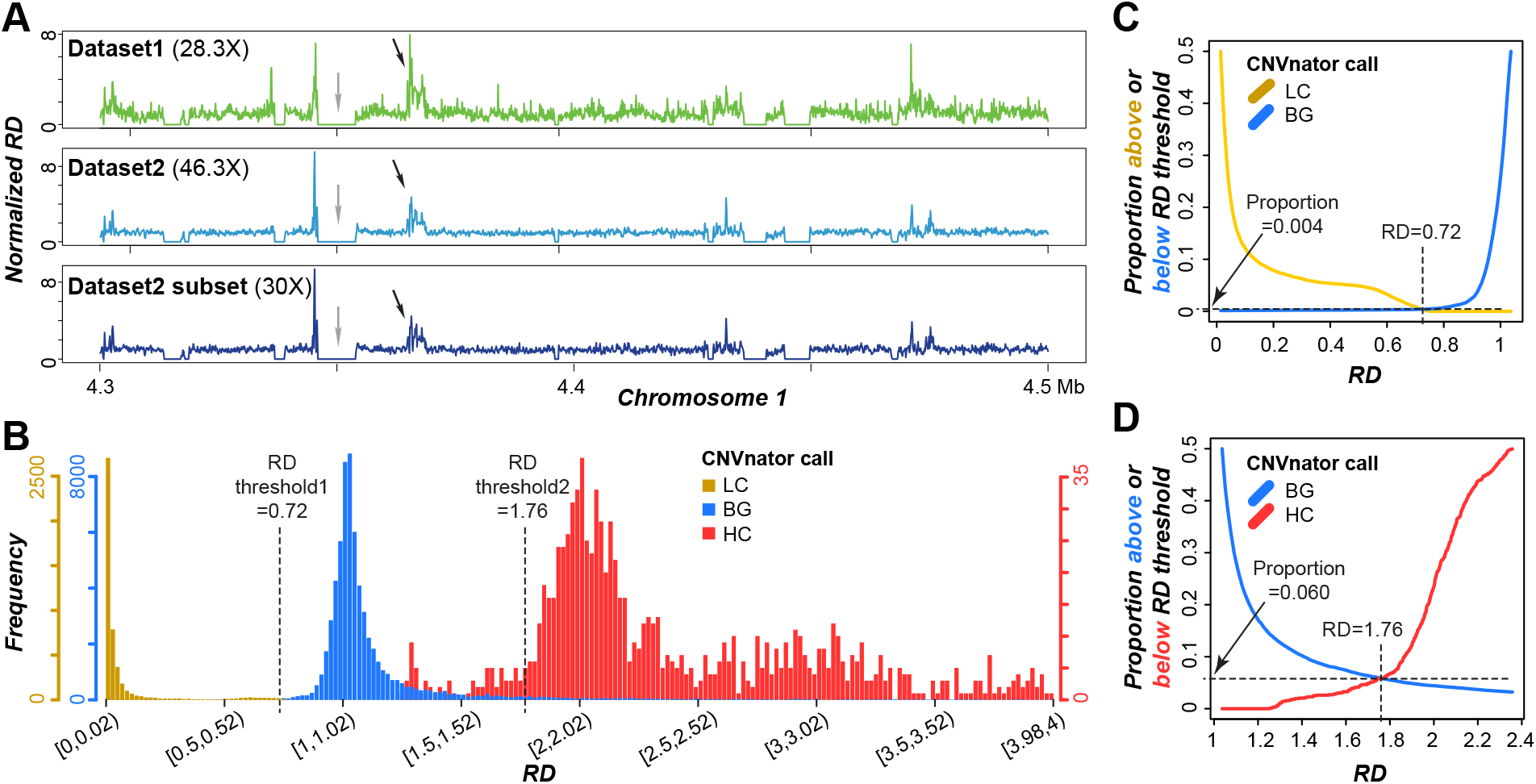
Identification of high, low, and background coverage regions. (**A**) Normalized read depth (RD) of 100bp bins using reads from dataset1, dataset2, and a dataset2 subset at 30-fold coverage in regions 4.3~4.5Mb on Chromosome 1. Black and grey arrows show regions with high read coverage and low coverage, respectively. (**B**) RD distributions of high coverage (HC), background (BG), and low coverage (LC) regions called by CNVnator. Dashed lines at RD=0.72 and 1.76: threshold values defining the boundaries between LC and BG regions and between BG and HC regions, respectively. (**C,D**) Proportion of called regions above or below RD value thresholds distinguishing LC and BG regions (**C**) and distinguishing BG and HC regions (**D**). Vertical dashed line: threshold value.

The comparably higher RD variance in dataset1 may be due to lower sequencing quality and shorter read lengths that contribute to erroneous read mapping and may lead to overestimates of HC and LC regions. Thus, only results based on dataset2 were discussed further. Based on the RD values, HC, LC, and BG regions were identified with CNVnator (see **Methods**). However, we found that RD distributions of HC, LC and BG regions called by CNVnator were overlapping (Fig. 1B). Given our goal is to identify HC and LC regions with high confidence, two threshold RD values were chosen: 0.72 and 1.76 that minimize the overlap between LC and BG regions (Fig. 1C), and that between BG and HC regions (Fig. 1D), respectively. Thus, HC regions were defined as regions with RD > 1.76, and LC regions were those with RD < 0.72. BG regions had RD values between 0.72 and 1.76. This resulted in 1,156 HC, 19,451 BG and 15,034 LC regions. The HC and LC regions account for 0.6% (5.1 Mb) and 9.7% (79.6 Mb) of the genome, respectively. The regions that are not classified as HC, LC, or BG due to this dual thresholding scheme are referred to as “other” regions. As expected, 95.5% of LC regions contain gaps filled with Ns, whereas only 18.2% of HC and 42% of BG contain Ns (**Fig. S1A**). In addition, the median lengths for LC, BG, and HC regions are 1.70, 16.30, and 2.80Kb, respectively (**Fig. S1B**).

### Prediction of HC, LC, and BG regions with a multi-class model using seven genomic features

After the HC/LC/BG regions were defined, we established machine learning model predicting which regions would be HC, LC, or BG regions. By predicting HC/LC/BG regions, we would have a better understanding of what the contributing genomic characteristics were, especially for HC regions where misassembly mostly likely have occurred. Starting out, we used seven features (referred to as base features, yellow box, Fig. 2A), including GC content, density values: all genes, tandem genes, non-tandem genes, pseudogenes, transposable elements (TEs), and simple sequence repeats (SSRs) of genomic regions to build a 3-class (HC, LC, or BG) model (referred to as Model 1, Fig. 2A, **Table S2**) using the Random Forest algorithm (33). We should emphasize that an independent, test set (10%) of HC, LC, and BG regions were set aside that was not used for model building. Thus, the test set was ideal for validating our models. Using Model 1, 61.9%, 83.9%, and 74.8% of HC, LC, and BG regions, respectively, were correctly predicted (Fig. 2B). Here the percentage true cases predicted correctly is defined as the recall value. Importantly, the testing set not used for model training were predicted with a similar recall (Fig. 2B). To jointly consider both recall and precision (% predictions that are correct), we determined the F1-score that is the harmonic mean of precision and recall. In our machine learning pipeline, we started with equal numbers of training and testing HC, LC, and BG regions (33% each). Thus, random guess would lead to an accuracy of 33% and F1=0.33. On the other hand, a perfect model would have an accuracy and an F1 of 1. Model 1’s F1=0.73 (Fig. 2A), while it was much better than random guess, the F1 was far from perfect.

**Figure 2.**
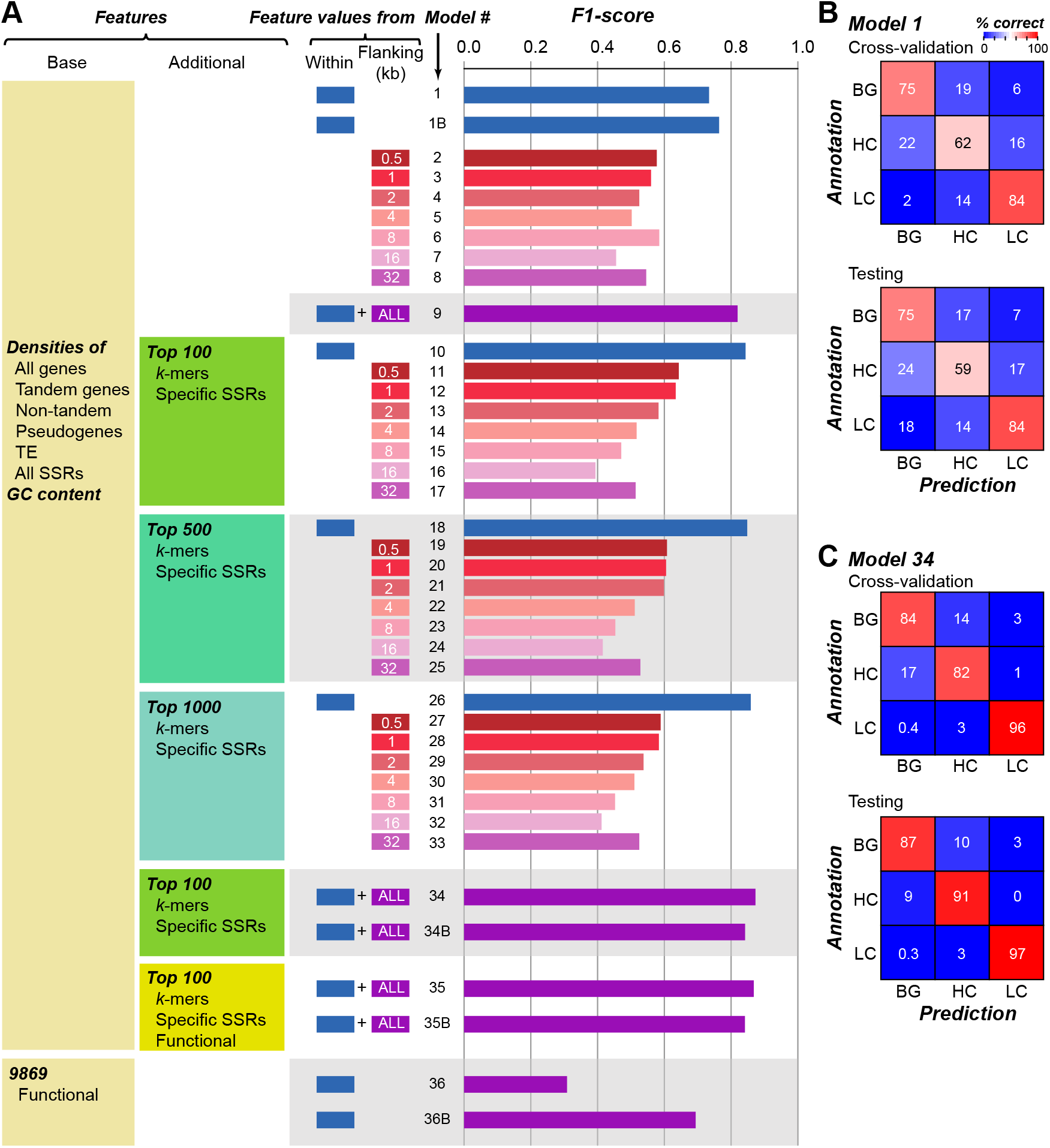
Prediction of HC/LC/BG regions using features of genomic sequences. (**A**) Average F1 of prediction models using different feature sets. The first two columns indicate feature sets used including Base features used in most models and additional features for model subsets. The third and fourth columns indicate whether feature values were derived from sequences within HC/LC/BG regions and/or flanking regions (0.5~32Kb), respectively. In the flanking region column, “all” refers to a combined set of values from all seven flanking regions. The fifth column contains model number (#), where B indicates binary model. (**B**) Confusing matrices of cross-validation and testing datasets based on Model 1. Color scale: % correct prediction. **(C)** Confusing matrices of cross-validation and testing datasets based on Model 34.

### Defining HC, LC, and BG regions with additional features

To improve upon Model 1, we included additional features from two sources. The first was the same seven base features but with values from flanking regions. The rationale was that the regions right next to HC, LC, and BG regions may have similar properties which can contributed to a better model. To assess this, we first build prediction models using only sequences flanking HC, LC, and BG regions by 0.5, 1, 2, 4, 8, 16, and 32kbs to build seven models (Model 2-8) and found that the performance of these models was worse than that of Model 1 (accuracy=46~58%, F1=0.46~0.58, Fig. 2A and **Table S2**). In addition, as the sizes of the flanking regions increased, the prediction performance decreased (Fig. 2A) This is likely because flanking regions can be of different types, i.e., a region flanking an HC region may be LC and/or BG regions. However, this is not because these regions are not important. When the features used for building Model 1 were combined with those for Model 2-8, the resulting model (Model 9) had a substantially improved F1=0.82 (Fig. 2A) compared to Model 1 (F1=0.74). This finding suggests that sequences flanking the HC/LC/BG regions, by themselves insufficient, have information that are useful for the prediction task.

In addition to flanking region, we focused on dissecting if HC, LC, and BG regions have different sequence composition—instead of compositions of much longer sequences (genes and transposons) or single nucleotides (GC content), we investigated whether specific SSRs (repeats with 2-64 bp units, 156,444 features) and/or *k*-mer (1-6bp, 5,460 features) may be prevalent in HC, LC, or BG regions. Because the number of these SRR and *k-*mer features was large, we first identified a subset of SSR and *k-*mer features with *p*-value < 0.05 (Kruskal-Wallis H test) among HC/LC/BG regions. Top 100 SSR and *k-*mers were further selected with a feature selection algorithm (see **Methods**). By incorporating these top SSR and *k-*mer features with seven base features to predict HC/LC/BG regions (Model 10), the performance of Model 10 (F1=0.84, Fig. 2A) was even better than the Model 9 (F1=0.82) that did not consider SSRs and *k-*mers but flanking regions. Thus, there exist substantial differences in the short sequence compositions among HC/LC/BG regions. We also included the top 500 and the top 1000 *k*-mers/SSRs to create Model 18 and Model 26 that improved performance further with F1=0.85 and 0.86, respectively (Fig. 2A). Although additional *k*-mers/SSRs may further improve predictions, they likely have diminishing contribution judging from the small F1 differences between Model 10, Model 18 and Model 26 (blue bars, Fig. 2A). Next we combined the features used in Model 9 and those from used in Model 10 to establish an all-inclusive Model 34 with 256 features that had F1=0.87 (Fig. 2A). Importantly, >84% BG, >82% HC and >96% LC regions are correctly predicted in both training and test (not used in model training) datasets (Fig. 2C).

### Features important for the prediction of HC/LC/BG regions

Model 34 has the highest F1=0.87 using 256 features (56 base features from 8 regions, 100 top *k-*mers, and 100 top SSRs, Fig. 2A). We next evaluated which features were among the most informative in distinguishing HC/LC/BG regions (highest feature importance values, see **Methods**). **Table S3** lists the importance values of all 256 features. We found that three types of features stand out: *k-*mers (median importance rank=57), GC contents (median rank=66), and density of TEs (median rank=70). TEs have long been implicated in their contributions to misassembly due to their lengths and high degree of similarities (3). Interestingly, the HC regions tend to have a significantly lower TE density compared to BG regions (Fig. 3A), likely reflecting genomic regions differing in recent transposition events. In contrast, LC regions have the highest TE density, although distribution of LC regions across the genome is not correlated with TE distribution (Spearman’s *rho* = 0.09, Fig. 3B). Furthermore, when we randomly reshuffled HC/LC/BG regions designations 1,000 times and determined the correlation distribution between prevalence of TEs and random genomic regions, the observed correlation value of LC regions was significantly lower than that of the random expectation (z-score=-3.0, Fig. 3C). One potential reason is that the assembler may be confused by the repetitive nature of TE and short length of sequencing reads to assemble sequences correctly, resulting in gaps filled with Ns in LC regions (**Fig. S1A**) with TE sequences at the breakpoints in one or both ends, which in turn led to higher TE density in LC regions.

**Figure 3.**
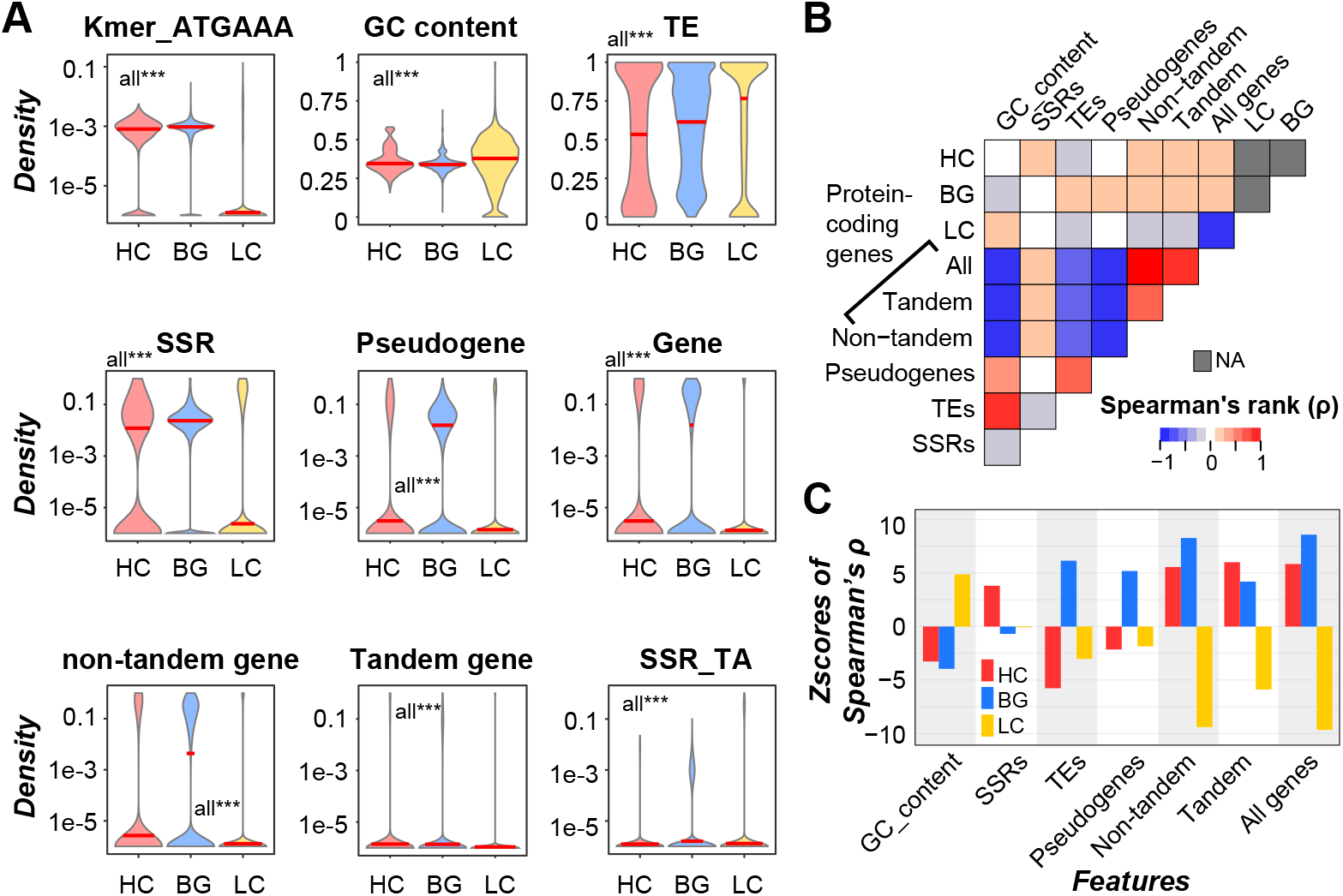
Relationships between genomic features and HC, LC and BG regions. **(A)** Differences in density distributions of important features between HC/LC/BG regions. ***: *p*<1e-3. (**B**) Correlation between densities of HC/LC/BG regions and other genome features across the genome in bins of 500Kb. Color scale: Spearman’s rank correlation coefficient (ρ). (**C**) Z-scores of observed ρ calculated using the distribution of 1000 random Spearman’s ρ as the null distribution. First, regions were selected so they were the same number and length as true HC/LC regions (not overlapping with each other). The remaining genomic regions not occupied by the selected random regions were taken as randomly expected BG regions. Then correlation between densities of randomly selected regions and a genomic feature was calculated to establish the null distribution.

As for GC content, it is well documented that, specifically for short read sequencing with the Illumina platform, GC-rich and GC-poor regions tend not to be sequenced and thus contribute to regions with low coverage or breakpoints in assemblies (4,37). Consistent with this, LC regions have significantly higher GC content compared to HC and BG regions (Fig. 3A). We also found that a major reason that the top 100 *k*-mers were important was because of their high AT content (88% with AT content ≥ 80% and 100% with AT content ≥ 60%, **Table S3**). In addition, the top two SSRs with highest importance in the prediction were “AT” (ranked 152) and its reverse complement “TA” (ranked 153). Contrary to densities of individual SSRs which generally had very low ranks (i.e., less important), densities of all SSRs ranked in the middle (108), indicating that it is more informative in distinguishing HC/LC/BG regions to consider SSRs as a whole. Consistent with this, when only the top 128 features were used, where the density of all SSRs was considered but no individual SSR feature, the model’s performance didn’t decline (Model 34_4 in **Table S2**).

Pseudogenes (median rank=131), all protein coding genes (139), non-tandem duplicates (141), and tandem duplicates (154) also ranked in the middle (**Table S3**). Densities of these genomic features in flanking regions ranked similarly or even higher than densities in HC/LC/BG regions (**Table S3**), suggesting the differences in genomic environment around HC/LC/BG regions. By determining the correlations between the prevalence of HC/LC/BG regions and the prevalence of genomic features in corresponding regions, we found that genomic regions with high densities of not only HC, but also BG regions tend to have higher gene density, regardless if tandem and non-tandem genes were separated or not (all ρ>0.12, *p-*values <2.0e-6, Fig. 3B and **Table S4**). Because LC regions tend to contain Ns, regions with higher LC density are expected to have lower gene density (all ρ< −0.17, *p*-values < 2.2e-11, Fig. 3B and **Table S4**). Given that density of protein coding genes is informative in distinguishing among HC/LC/BG regions, we next asked whether the types of genes, in terms of functional aspects (e.g., Pfam domains, biological processes and metabolic pathways) also impact RD. To test this, top 100 functional characteristics (domains, biological processes and pathways) of genes within HC/LC/BG regions (9869 features in total) were also combined with Model 34 to build Model 35 (Fig. 2A, **Table S2**). However, there was no apparent improvement compared to model 34 (model 35’s overall accuracy = 87%, F1 = 0.87).

### Features important in binary classification model distinguishing HC and BG regions

Although the importance analysis allows us to pinpoint the features crucial for Model 34’s performance, Model 34 is a 3-class model and thus it is not straightforward to tell if a feature is important because it allows us to distinguish HC from BG and LC regions or other scenarios. Because we are mostly interested in assessing why HC regions exist, we next establish a binary classification model to distinguish HC from background regions. Using the same features as in Model 1, we established a new Model 1B (B=binary) with HC and BG regions as classes and found that it has an accuracy=84% and F1=0.76 (Fig.2A). As expected, Model 34B that used the same feature set as Model 34 for binary classification had even better accuracy=92% and F1=0.84 (Fig.2A). Like Model 35 which didn’t lead to improved classification among HC/LC/BG regions by including functional features (**TableS2**), the comparable binary Model 35B had the same performance as Model 34B (Fig.2A, **Table S2**). However, models with only functional features had accuracy=58% and F1=0.69, which is much better than random guess (**TableS2**), suggesting that functional features are still informative in distinguishing HC from BG regions.

As expected, the important features for binary classification of HC and BG regions differ from those for the 3-class model. For example, densities of *k-*mer in Model 34B (median rank=73) and in Model 35B (median rank=72) were no longer the most important feature categories as in Model 34 (median rank=57) and 35 (median rank=55, **TableS3,5,6,7**). In contrast, GC content, density of TE and SSRs had the highest median ranks (12, 13, and 57 in Model 35B, respectively). For HC regions, one hypothesis for their presence is due to the presence of multiple copies of highly similar sequences arranged in tandem that are misassembled. If this is true, one would expect that SSRs and tandem genes would tend to be co-localized with HC regions compared to BG regions. Consistent with the above hypothesis, although the density of SSRs in HC regions was slightly lower than BG regions (Fig. 3A), it was significantly higher than randomly expected (z-score=3.81, Fig. 3C). In contrast, density of SSRs in BG regions was slightly lower than random expectation (z-score=-0.69 Fig. 3C). In addition, the density of SSRs across genome is positively correlated with HC, not BG, regions (Fig. 3B), and the flanking regions of HC also have higher density of SSRs than those of BG regions (**Fig. S2**). These results suggest the potential contribution of SSRs to misassembly in HC regions, which resulted in underestimation of SSRs density in HC regions. The situation is similar for tandem genes, although it is not as important as SSRs (median rank=155). The observed correlation value (ρ) for HC regions was significantly higher than random expectation (z-score=6.0) compared to that for BG regions (z-score=4.1, Fig. 3C). Note that, although both have positive z-scores due to consideration of LC regions also, the higher z-score for HC regions indicates that tandem gene density is more prevalent in HC than in BG regions. Conversely, compared to BG regions, HC regions tend to have fewer non-tandem genes (Fig. 3C). Thus, the presence of tandem genes also contributes to misassembly.

### Properties of genes located in HC regions

In earlier section, functional characteristics (domains, biological processes and pathways) of genes within HC/LC/BG regions were also combined with the seven base features to build Model 35 and 35B (**Table S2**) that resulted in no apparent improvement compared to model 34 and 34B that did not incorporate functional characteristics (Fig. 2A). This may be because properties contributing to the enriched presence of genes with certain functional characteristics were already considered, it is also possible that, due to the large number of features considered and the fact that functional characteristics tend to be lower ranked, the contribution of functional characteristics was not apparent in Model 35 and 35B because other features dominated. To assess the extent to which functional characteristics could be used to predict whether a genomic regions would be BG, HC, or LC, we established three-class models using only functional features and found that it had accuracy=41% and F1=0.31, very close to random guess, no matter how many features were selected (Model 36, Fig. 2A). However, binary model for classifying HC and BG regions using only functional features had accuracy=57.9% and F1=0.69, indicating that they were informative (Model 36B, Fig. 2A, **Table S2**).

To assess what types of genes tend to be located in BG and HC regions, we first determined if the numbers of different types of genes (**Table S8**) were over or under-represented in HC compared to BG regions. By generating 10,000 datasets with randomized HC locations, we established the randomly expected numbers of different gene types and the resulting null distributions were used to assess the statistical significance of observed numbers of different gene types (Fig. 4A). In this analysis, two types of genes stand out, specialized metabolism (SM) protein coding genes and RNA genes. SM genes has a z-score=2.1, indicating that SM genes tend to be found in HC regions and thus misassembled. This is consistent with the findings that SM genes tend to belong to large gene families, located in tandem clusters, and be recently duplicated (19,20). However, genes in larger families are not necessarily in HC regions (black arrow, Fig. 4A) and number of SM genes that are tandemly duplicated is not significantly higher than random expectations (green arrow, Fig. 4A). Thus, it is likely that the over-representation of SM genes in HC regions is due to their higher duplicate rate, but not always through tandem duplication, resulting in closely related copies that were misassembled. It also can be because tandem duplicated SM genes were misassembled together, which makes the number of tandem duplicated SM genes underestimated. In addition to SM genes, surprisingly, non-coding RNAs (ncRNAs) tend to be enriched within HC regions (z-score=2.5, purple arrow, Fig. 4A). We speculate that tomato ncRNA regions may have a higher than average rate of recent duplications, which would indicate there are more ncRNA regions than annotated and ncRNA expression levels may be overestimated because multiple ncRNA regions are assembled together.

**Figure 4.**
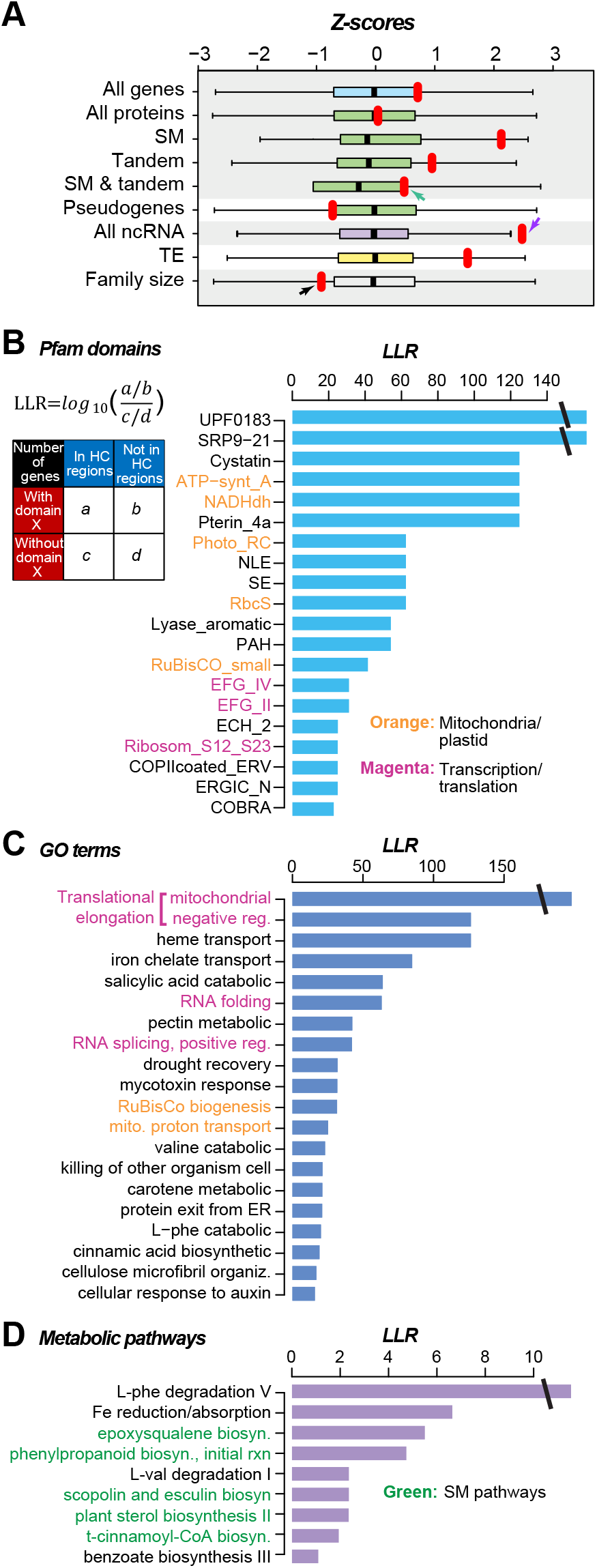
Sequences found in HC regions and their functions. (**A**) Z-score calculated by comparing observed total number of features in true HC regions (red dots) to normalized distributions (boxplots) of the numbers of different genomic features overlapped with 10,000 randomly sampled BG regions (based on the number and length distribution of HC regions). (**B**) Top 20 Pfam domains with highest Log Likelihood Ratio (LLR, see upper-left insert and **Methods**), indicating enrichment within HC regions. (**C**) Top 20 GO terms with highest LLR. (**D**) 9 metabolic pathways with LLR > 1. Orange and magenta fonts indicate mitochondria/plastid and transcription/translation related processes, respectively, and green font shows specialized metabolism pathway.

Our finding that SM genes tend to be over-represented in HC regions suggests that genes with other functions may have similar behaviors. To address this, we asked if there was enrichment of any Pfam domain family, Gene Ontology biological process category, or TomatoCyc pathways. Given the number of domain families (**Table S9**), categories (**Table S10**) and pathways (**Table S11**) were large, multiple testing correction was applied and resulted in only one statistically significantly enriched entry (salicylic acid catabolic process). To assess if there are general patterns we may have missed due to the stringency of the multiple testing corrections, we examined the Log Likelihood Ratio (LLRs, see **Methods**) between the numbers of genes with or without a protein domain X and the numbers of genes within or out of HC regions (inserted table, Fig. 4B). Similarly, we examined the LLRs for biological processes (Fig. 4C) and pathways (Fig. 4D).

There are three general patterns that emerge. The first is the prevalence of nuclear encoded proteins responsible for mitochondrial and plastid functions among the Pfam domains and the GO categories with the highest LLRs—including ATP-synt_A: ATP synthase A chain, NADHdh: NADH dehydrogenase, Photo_RC: photosynthesis reaction center, and RbcS and RuBisCO_small: Ribulose-1,5-bisphosphate carboxylase small subunit (Fig. 4B), as well as mitochondrial proton transport and RuBisCo biogenesis (Fig. 4C). The second general pattern is the occurrence of domain/process related to transcription and translations—including various translational elongation factor G (EFG) domains, translational elongation-related functions, ribosomal proteins, RNA splicing (Fig. 4B,C). One outstanding property of genes that fit these two general patterns is their extremely high level of expression. Such high level of expression is known to lead to the generation of retrogenes and retro-pseudogenes with highly similar sequences that littered around various parts of the genomes (38,39) thus higher coverage within genomes. Consistent with this hypothesis, the average number of introns in genes found in HC regions was significantly lower than that in BG regions (2.13 vs. 4.38 in average, Kolmogorov-Smirov test, *p*=4.3e-09). The third general pattern is revealed from the few metabolic pathways with LLR value > 1 where five out of nine were SM pathways (Fig. 4D), as expected.

### Evaluation of HC region misassembly by comparing Short-read and Long-read assemblies

To assess the extent to which HC regions tend to be mis-assembled, Short-read assembly (query) was aligned to Long-read assembly (subject) using MUMmer (36) (see **Methods**), and the aligned regions were shown in **Table S12**. Aligned regions were classified into six categories (see **Methods**, Fig. 5A): 1) non-duplicated, <u>C</u>orrectly assembled (C1, 681.0 Mb); 2) non-duplicated, <u>M</u>is-assembled (M1, 0.6 Mb); 3) locally duplicated (i.e., on the same chromosome of the Long-read assembly), correctly assembled (C2, 11.5 Mb), 4) locally duplicated, mis-assembled (M2, 8.2 Mb); 5) non-locally (on different chromosome) duplicated, correctly assembled (C3, 29.5 Mb); and 6) non-locally duplicated, mis-assembled (M3, 4.9 Mb). We found that 86.3Mb of Short-read assembly regions, mostly consisted of Ns, that could not be aligned to the Long-read assembly. Since LC regions tend to consist of Ns, it is not surprising that 93% of the total length of LC regions had no match in Long-read assembly (Fig. 5B). Thus, LC regions were not further examined. Of the 5.1 Mb HC regions, 0.03 Mb (0.53%), 0.97 Mb (19.1%) and 0.44 Mb (8.7%) were in misassembled categories M1, M2 and M3, respectively (Fig. 5B). Compared to HC regions, the proportion of misassembled regions in BG were significantly lower (0.08 %, 0.9% and 0.5% for M1, M2, and M3, respectively; Fisher’s Exact tests, all *p<*2.8e-09). Among the three categories of misassembled regions, only 1.9% HC regions (M1) did not have duplicated subject regions in the Long-read assembly. The majority of misassembly (67.4% and 30.8%) was due to duplications (M2 and M3, respectively), suggesting that HC regions were much more likely to be mis-assembled due to duplications, especially when duplications occurred on the same chromosome.

**Figure 5.**
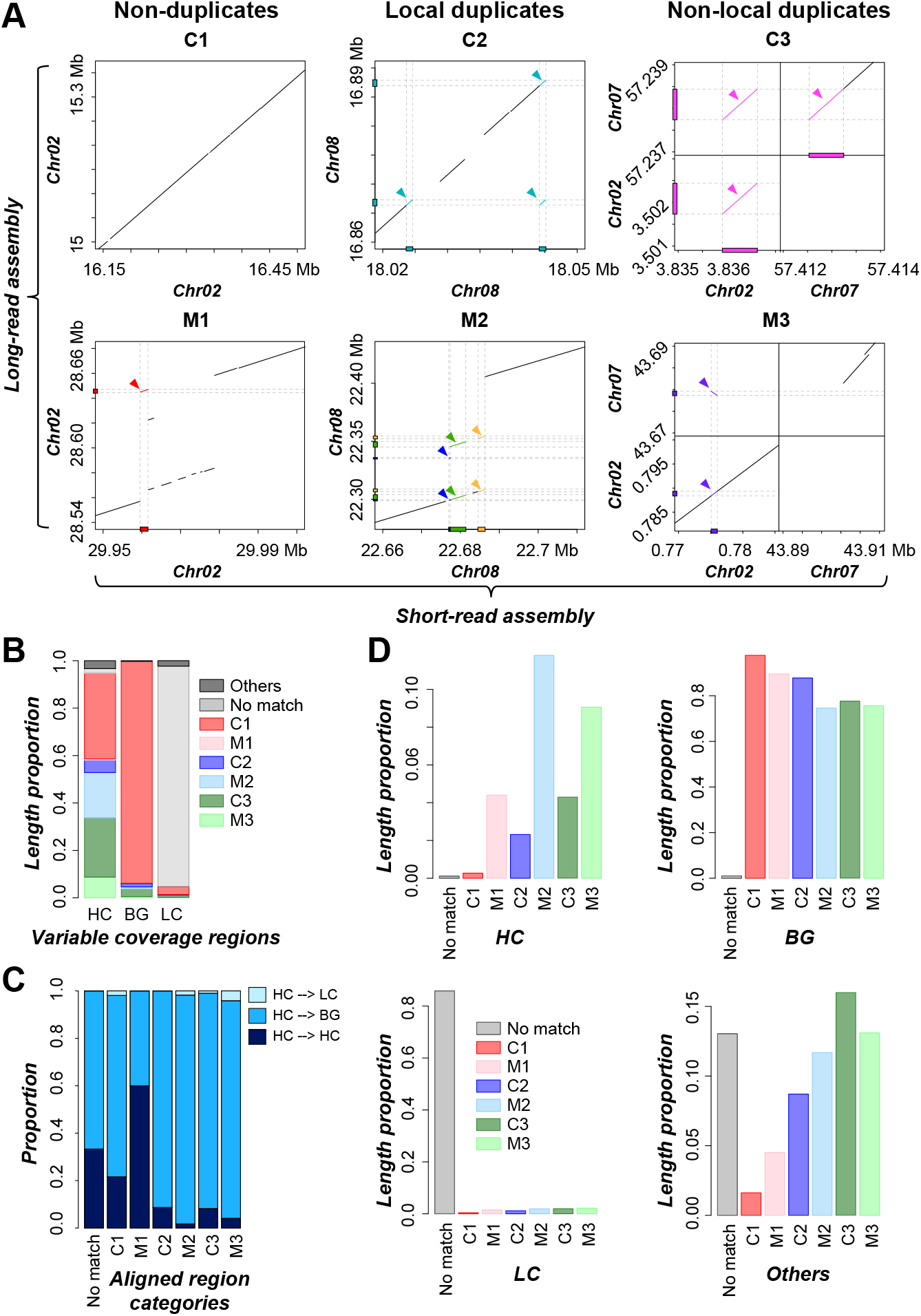
Categories of correctly and mis-assembled regions based on alignments of Short- and Long-read assemblies. (**A**) Dotplots of example genomic regions of six categories of regions aligned between Short- and Long-read assembly. Color boxes and lines: corresponding regions between assemblies bound by the dotted lines. (**B**) Proportion of total length of HC, BG, or LC regions in each of six aligned region categories in (**A**). (**C**) Proportion of the total number of Short-read assembly HC regions defined as HC, LC, BG regions in Long-read assembly for each aligned region category. (**D**) Proportion of the total length of an aligned region category overlapped with HC, LC, BG, or other regions. For each aligned region category, the values from HC, BG, LC, and other add up to be 1.0.

Thus far, HC regions were defined by mapping short reads to the Short-read assembly. To evaluate whether these Short-read assembly-based HC regions were still classified as HC in the Long-read assembly, the shot reads were also mapped to the Long-read assembly to determine variable coverage regions. This resulted in 297 HC, 4,479 BG and 1,971 LC regions based on the Long-read assembly. Importantly, among 1,156 Short-read assembly-based HC regions, once we map the reads to the Long-read assembly, only 88 (7.6%) overlapped with the Long-read assembly-based HC regions. In addition, among mis-assembled HC regions (coverage defined using the Short-read assembly), 96.5% and 91.8% of M2 and M3 were identified as BG based on the Long-read assembly (Fig. 5C, **Table S15**). These findings further suggest that higher than usual read coverage is a good indicator of misassembly.

HC regions tend to be misassembled compared to BG or LC regions (Fig. 5B). If we broke down the six aligned region categories (Fig. 5A), it was clear that HC regions have higher proportion of M2 (11.8%) and M3 (9.1%) compared to other categories (0.3-4.4%, Fig. 5D). Nonetheless, 74.6% of M2 and 75.7% of M3 were identified in BG regions (Fig. 5D). One potential reason is that some true HC regions were identified as BG in our analysis. If that was the case, we would expect misassembled BG regions (which presumably were HC) would have significantly higher read coverage compared to correctly assembled BG regions. Consistent with this expectation, the median RDs of 100bp BG region bins that were misassembled (1.13 and 1.28 for M2 and M3, respectively) were higher than the median RDs of correctly assembled BG bins (1.03 and 1.06 for C2 and C3, respectively; Wilcoxon signed-rank tests, both *p<*2.2e-16). In addition to read coverage differences, we found that misassembled BG regions tend to be much shorter (median lengths=698bp for M2/M3 combined) than misassembled HC regions (2,328bp; **Fig. S1C**; Wilcoxon signed-rank test, *p* < 2.2e-16). This is likely because CNVnator (21) merges adjacent bins based on read depth similarity and in doing so, shorter regions with variable coverage may not be identified. In any case, the read coverage difference is small. Thus, if we relaxed the HC detection threshold, it would significantly increase the false HC calls by calling true BG regions as HC.

### Genome features distinguishing correctly and mistakenly assembled regions

To understand why some HC regions were not identified as mis-assembled, using the same genome features as in Model 35 for classifying HC, BG, and LC regions, a binary classification model (Model 37) was built to distinguish HC regions consisted of mainly M2/M3 (>50%, referred to as the HC_M2/M3 class) or mainly C2/C3 (>50%, referred to as the HC_C2/C3 class). Model 37 resulted in an F1 = 0.79 (balanced positive and negative classes, thus the background F1 was 0.5), indicating that mis- and correctly assembled HC regions were significantly distinct from each other in certain genome features. Among the top 20 most important features (Fig. 6A, detailed distribution of feature values was shown in **Fig. S4**), interestingly the most informative ones were those of regions flanking the misassembled regions. The flanking sequences of HC_M2/M3 regions tend to have higher densities of SSR, pseudogenes, TE, and non-tandem genes. At first it was surprising that it was the features in the flanking regions that were informative. In hindsight, if a region was misassembled, the distinguishing signature would likely be buried with it. From the flanking regions, one can better defined whether the sequence in between is problematic, i.e., in our case, misassembled. Within HC_M2/M3 regions, there tend to be higher densities of four types of *k*-mers including TATTTC, TGTAA, ATACTT, and GATTTT. However, it is not clear why these *k*-mers are informative.

**Figure 6.**
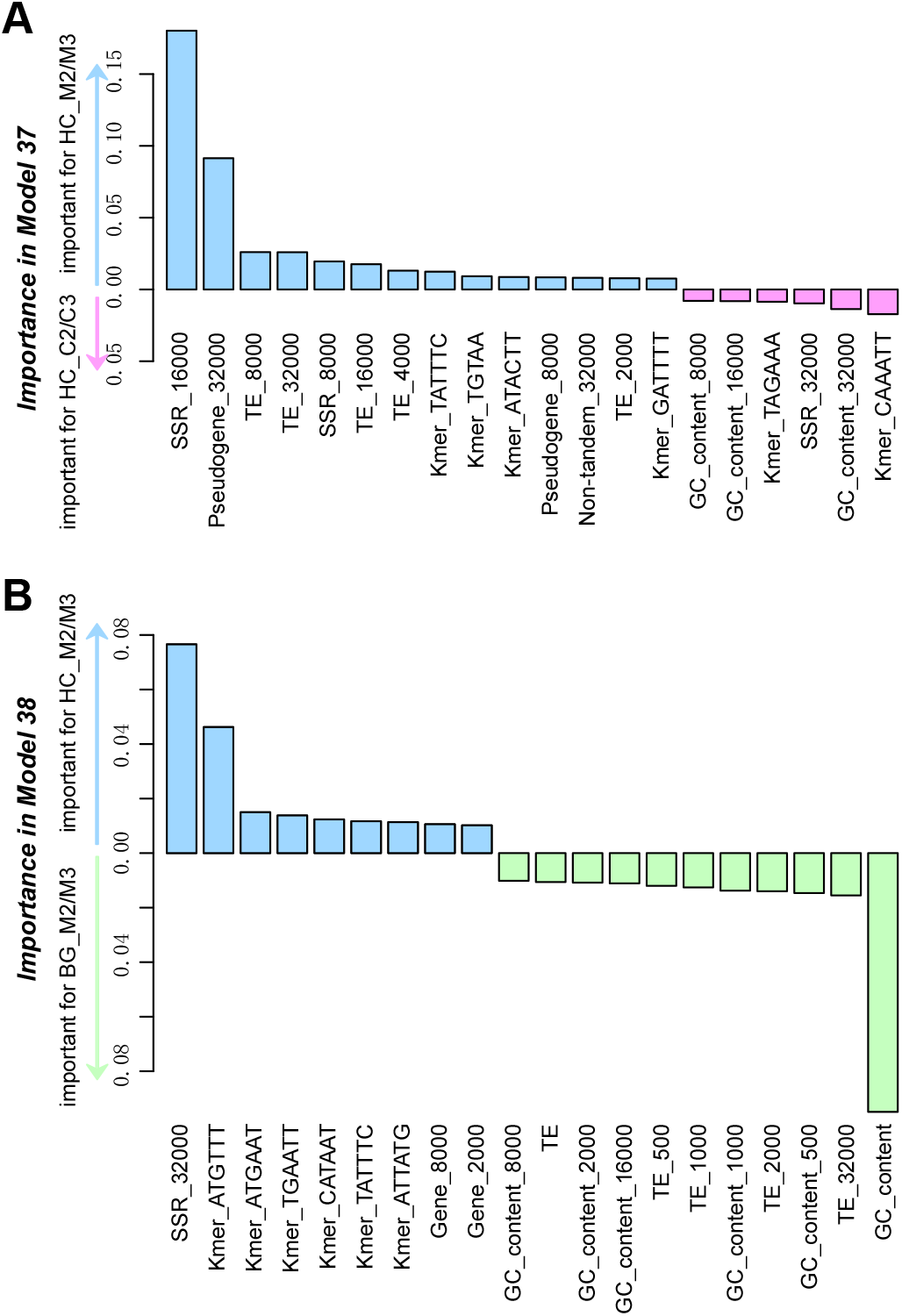
Important features in Model 37 and Model 38. **(A)** Model 37 is for classifying HC_M2/M3 and HC_C2/C3 to assess the features important for predicting misassembled HC regions. **(B)** Mode 38 is for classifying HC_M2/M3 and BG_M2/M3 to assess if misassembled HC and BG regions have distinguishable features. Bar height: importance value in the Random Forest model. Blue, red, green: median values of features are higher in HC_M2/M3, HC_C2/C3, and BG_M2/M3, respectively. The distributions of feature values were shown in **Fig. S4** and **Fig. S5** for Model 37 and Model 38, respectively.

In the above modeling exercise, we were able to distinguish HC regions that were likely misassembled from those that were not. Recall that not only HC regions contain misassembled sequences, in fact BG regions have higher proportion of M2/M3 regions (Fig. 5D). To further dissect their differences and to understand why some mis-assembled regions were not detected as HC, another model (Model 38) was built to distinguish HC_M2/M3 and BG_M2/M3 (>50% of a HC or BG region overlapped with M2/M3), using same features as in Model 35. The resulting F1 was 0.77. Among the top 20 most important features, BG_M2/M3 regions tend to have higher GC content (41.3%) and TE density (0.92) compared to HC_M2/M3 regions (GC=33.6%, TE density=0.61, Fig. 6B, feature value distributions shown in **Fig. S5**). This trend is also true when comparing BG_M2/M3 and HC_M2/M3 flanking regions (Fig. 6B, **Fig. S5**). Interestingly, the comparatively lower GC content in HC_M2/M3 regions (33.6%) is more similar to the 36.6% overall GC content in the tomato genome. In addition, the GC content in genic region is at 42.4%, suggesting BG_M2/M3 and HC_M2/M3 may be located in relatively gene-rich and poor regions, respectively. Contrary to this expectation, however, HC_M2/M3 regions tend to have significantly higher gene density (average=0.16, rank=142) compared to BG_M2/M3 regions (average=0.04, Wilcoxon signed-rank test, *p*=1.3e-15). This is also true when comparing flanking regions (Fig. 6B and **Fig. S5**). With regard to TE, we have already shown that HC regions tend to have a significantly lower TE density compared to BG regions regardless whether they are misassembled or not (Fig. 3A). Taken together, these predictive models perform well for distinguishing mis-from correctly assembled HC regions and for predicting whether a misassembled region lies in BG or HC regions. Using model interpretation strategies, we are able to identify salient genome features underlying the models’ ability to make good quality predictions.

## CONCLUSION

Although the third-generation sequencing such as the PacBio (7) and 10X (8) are now available, the majority of existing genome assemblies were derived from short-read based technologies. With the goal of evaluating genome assembly quality by assessing short read coverage distribution, we identified 1,156 HC and 15,034 LC regions in tomato genome assembly SL2.50 (24). These variable coverage regions collectively accounted for ~10% of the genome assembly, indicating the severity of the issue. By applying machine learning methods, we found that HC and LC regions can be predicted with high accuracy. High GC content and TE density are the major factors contributing to the low read coverage or the break point of assembly, while SSRs and tandem duplicates, especially specialized metabolism genes, tend to be in HC regions, potentially leading to mis-assembly due to high sequence similarities. By comparing Short- and Long-read assemblies, 27.8% of HC regions were potentially mis-assembled due to duplications. In addition, 91.8% of misassembled HC regions no longer defined as high coverage when we mapped the short reads to the Long-read assembly. Our results highlight the extent to which variable coverage in a Short-read assembly contribute to misassembly, particularly when they are flanked by TEs and tandemly duplicated sequences.

Misassembled regions that are duplicated were detected in both HC and BG regions. It is straightforward to appreciate why misassembled HC region strongly correlated with duplication in the Long-read assembly—higher read coverage is a strong indication that more than one genome regions are likely assembled together. However, it is not as obvious why BG regions would be duplicated. There are four explanations. First, HC regions could be underestimated in our approach. Misassembled BG regions tend to have slightly higher read coverages compared to correctly assembled ones. Second, related to the first explanation, after the partitioning of the genome to HC/LC/BG regions, read depth varies continuously across the genome, and there are no sharp boundaries between HC and BG regions (as opposed to between LC and BG), we established a threshold to define HC and BG regions. As a result, regions with coverage near the defined threshold may be mis-labeled. Third, we defined a genomic region into six categories based on whether it is misassembled or not, duplicated or not, and, if duplicated, locally or not. This analysis is based of anchored matches of the Short and Long-read assemblies and thus alignment methods and their parameter choice is expected to impact our findings. Finally, the current tomato Long-read assembly still has scaffolds that cannot be mapped to any chromosomes, which may also contribute to an underestimate of misassembled regions using read coverage. Although there remain areas for further improvement, our results highlight the utility of detecting HC regions in short-read based assemblies for identifying potential mis-assembled regions. Although not all HC regions have evidence of misassembly based on the Long-read assembly, we showed that, with the machine learning model, misassembled HC regions can be readily distinguished from those that are correctly assembled.

Unlike methods developed for evaluating genome assembly continuity, like LTR Assembly Index (15) and MaGuS (14), here we focused on identifying misassembled regions based on variation in read coverage across the genome, and uncovering the underlying contributors in genome sequences using machine learning. Using tools for identifying regions with significantly high or low read coverages in estimating CNVs among individuals (21–23) and comparison to Long-read assembly, we discovered potential misassembled genomic regions. Through machine learning, our study combined a large number of genomic and functional annotation features. The resulting model provides an extensive, quantitative estimate of our current state of understanding of factors contributing to variable genome coverage and assembly issues in short read assemblies. These variable coverage regions account for ~10% of tomato genomes. In addition, HC regions tend to be misassembled. Considering that the presence of mis-assembled regions can impact genome-wide studies significantly, their detection prior to genome-wide analysis should be conducted to reduce the impact of misassembly due to variable coverage.

## FUNDING

This work was partly supported by the National Science Foundation [IOS-1546617, DEB-1655386 to S.-H.S.]; and U.S. Department of Energy Great Lakes Bioenergy Research Center [BER DE-SC0018409 to S.-H.S.].

## ACKNOWLEDGEMENTS

We thank the two anonymous reviewers whose comments/suggestions helped improve and clarify this manuscript, and thank Melissa Lehti-Shiu, Christina Azodi and Siobhan Cusack for discussion.

## COMPUTATIONAL RESOURCES

All the scripts used in this study are available on Github at: https://github.com/peipeiwang6/Evaluating-misassembly/

## Notes

#### Summary of Updates

Based on a recently available PacBio Long-read assembly, we have conducted three new analyses: (1) comparing the Short and Long-read assemblies to identify potentially mis-assembled regions, (2) evaluating the overlaps between HC/LC/BG regions and misassembled regions, (3) establishing new machine learning models distinguishing correctly and mistakenly assembled HC regions, and (4) establishing new machine learning model classifying misassembled HC and BG regions.

## REFERENCES

1. Heather,J.M. and Chain,B. (2016) The sequence of sequencers: The history of sequencing DNA. Genomics, 107, 1–8.

2. Consortium,I.H.G.S. and International Human Genome Sequencing Consortium (2001) Erratum: correction: Initial sequencing and analysis of the human genome. Nature, 412, 565–566.

3. Treangen,T.J. and Salzberg,S.L. (2011) Repetitive DNA and next-generation sequencing: computational challenges and solutions. Nat. Rev. Genet., 13, 36–46.

4. Chen,Y.-C., Liu,T., Yu,C.-H., Chiang,T.-Y. and Hwang,C.-C. (2013) Effects of GC bias in next-generation-sequencing data on de novo genome assembly. PLoS One, 8, e62856.

5. Aird,D., Ross,M.G., Chen,W.-S., Danielsson,M., Fennell,T., Russ,C., Jaffe,D.B., Nusbaum,C. and Gnirke,A. (2011) Analyzing and minimizing PCR amplification bias in Illumina sequencing libraries. Genome Biol., 12, R18.

6. Clavijo,B.J., Venturini,L., Schudoma,C., Accinelli,G.G., Kaithakottil,G., Wright,J., Borrill,P., Kettleborough,G., Heavens,D., Chapman,H., et al. (2017) An improved assembly and annotation of the allohexaploid wheat genome identifies complete families of agronomic genes and provides genomic evidence for chromosomal translocations. Genome Res., 27, 885–896.

7. Eid, J., Fehr, A., Gray, J., Luong, K., Lyle, J., Otto, G., Peluso, P., Rank, D., Baybayan, P., Bettman, B. et al. (2009) Real-time DNA sequencing from single polymerase molecules. Science, 323, 133–138.

8. Jain, M., Olsen, H.E., Paten, B. and Akeson, M. (2016) The Oxford Nanopore MinION: delivery of nanopore sequencing to the genomics community. Genome Biol., 17, 239.

9. Watson,M. and Warr,A. (2019) Errors in long-read assemblies can critically affect protein prediction. Nat. Biotechnol., 37, 124–126.

10. Sedlazeck,F.J., Lee,H., Darby,C.A. and Schatz,M.C. (2018) Piercing the dark matter: bioinformatics of long-range sequencing and mapping. Nat. Rev. Genet., 19, 329–346.

11. Zhuang,W., Chen,H., Yang,M., Wang,J., Pandey,M.K., Zhang,C., Chang,W.-C., Zhang,L., Zhang,X., Tang,R., et al. (2019) The genome of cultivated peanut provides insight into legume karyotypes, polyploid evolution and crop domestication. Nat. Genet., 51, 865–876.

12. Bertioli,D.J., Jenkins,J., Clevenger,J., Dudchenko,O., Gao,D., Seijo,G., Leal-Bertioli,S.C.M., Ren,L., Farmer,A.D., Pandey,M.K., et al. (2019) The genome sequence of segmental allotetraploid peanut Arachis hypogaea. Nat. Genet., 51, 877–884.

13. Edger,P.P., Poorten,T.J., VanBuren,R., Hardigan,M.A., Colle,M., McKain,M.R., Smith,R.D., Teresi,S.J., Nelson,A.D.L., Wai,C.M., et al. (2019) Origin and evolution of the octoploid strawberry genome. Nat. Genet., 51, 541–547.

14. Madoui,M.-A., Dossat,C., d’Agata,L., van Oeveren,J., van der Vossen,E. and Aury,J.-M. (2016) MaGuS: a tool for quality assessment and scaffolding of genome assemblies with Whole Genome Profiling^TM^ Data. BMC Bioinformatics, 17.

15. Ou,S., Chen,J. and Jiang,N. (2018) Assessing genome assembly quality using the LTR Assembly Index (LAI). Nucleic Acids Res., 46, e126.

16. Yang,L.-A., Chang,Y.-J., Chen,S.-H., Lin,C.-Y. and Ho,J.-M. (2019) SQUAT: a Sequencing Quality Assessment Tool for data quality assessments of genome assemblies. BMC Genomics, 19, 238.

17. Kozarewa,I., Ning,Z., Quail,M.A., Sanders,M.J., Berriman,M. and Turner,D.J. (2009) Amplification-free Illumina sequencing-library preparation facilitates improved mapping and assembly of (G+C)-biased genomes. Nat. Methods, 6, 291–295.

18. Vukašinović,N., Cvrčková,F., Eliáš,M., Cole,R., Fowler,J.E., Žárský,V. and Synek,L. (2014) Dissecting a hidden gene duplication: the Arabidopsis thaliana SEC10 locus. PLoS One, 9, e94077.

19. Chae,L., Kim,T., Nilo-Poyanco,R. and Rhee,S.Y. (2014) Genomic signatures of specialized metabolism in plants. Science, 344, 510–513.

20. Moore,B.M., Wang,P., Fan,P., Leong,B., Schenck,C.A., Lloyd,J.P., Lehti-Shiu,M.D., Last,R.L., Pichersky,E. and Shiu,S.-H. (2019) Robust predictions of specialized metabolism genes through machine learning. Proc. Natl. Acad. Sci. U. S. A., 116, 2344–2353.

21. Abyzov,A., Urban,A.E., Snyder,M. and Gerstein,M. (2011) CNVnator: an approach to discover, genotype, and characterize typical and atypical CNVs from family and population genome sequencing. Genome Res., 21, 974–984.

22. Miller,C.A., Hampton,O., Coarfa,C. and Milosavljevic,A. (2011) ReadDepth: a parallel R package for detecting copy number alterations from short sequencing reads. PLoS One, 6, e16327.

23. Li,S., Dou,X., Gao,R., Ge,X., Qian,M. and Wan,L. (2018) A remark on copy number variation detection methods. PLoS One, 13, e0196226.

24. Tomato Genome Consortium (2012) The tomato genome sequence provides insights into fleshy fruit evolution. Nature, 485, 635–641.

25. Ezura,H., Ariizumi,T., Garcia-Mas,J. and Rose,J. (2016) Functional Genomics and Biotechnology in Solanaceae and Cucurbitaceae Crops Springer.

26. Li,H. and Durbin,R. (2010) Fast and accurate long-read alignment with Burrows-Wheeler transform. Bioinformatics, 26, 589–595.

27. Wang,P., Moore,B.M., Panchy,N.L., Meng,F., Lehti-Shiu,M.D. and Shiu,S.-H. (2018) Factors Influencing Gene Family Size Variation Among Related Species in a Plant Family, Solanaceae. Genome Biol. Evol, 10, 2596–2613.

28. Benson,G. (1999) Tandem repeats finder: a program to analyze DNA sequences. Nucleic Acids Research, 27, 573–580.

29. Wang,Y., Li,J. and Paterson,A.H. (2013) MCScanX-transposed: detecting transposed gene duplications based on multiple colinearity scans. Bioinformatics, 29, 1458–1460.

30. Conesa,A. and Götz,S. (2008) Blast2GO: A comprehensive suite for functional analysis in plant genomics. Int. J. Plant Genomics, 2008, 619832.

31. Camacho,C., Coulouris,G., Avagyan,V., Ma,N., Papadopoulos,J., Bealer,K. and Madden,T.L. (2009) BLAST+: architecture and applications. BMC Bioinformatics, 10, 421.

32. Benjamini,Y. and Hochberg,Y. (1995) Controlling the False Discovery Rate: A Practical and Powerful Approach to Multiple Testing. J. R. Stat. Soc. Series B Stat. Methodol, 57, 289–300.

33. Breiman,L. (2001) Random Forests. Mach. Learn., 1, 5–32.

34. Pedregosa,F., Varoquaux,G., Gramfort,A., Michel,V., Thirion,B., Grisel,O., Blondel,M., Prettenhofer,P., Weiss,R., Dubourg,V., et al. (2011) Scikit-learn: Machine Learning in Python. J. Mach. Learn. Res., 12, 2825–2830.

35. Jones E, Oliphant E and Peterson,P. (2001) SciPy: Open Source Scientific Tools for Python. Available at: http://www.scipy.org/. (Accessed: 21st August 2019)

36. Kurtz, S., Phillippy, A., Delcher, A.L., Smoot, M., Shumway, M., Antonescu, C. and Salzberg, S.L. (2004) Versatile and open software for comparing large genomes. Genome Biol., 5, R12.

37. Dohm, J.C., Lottaz, C., Borodina, T. and Himmelbauer, H. (2008) Substantial biases in ultra-short read data sets from high-throughput DNA sequencing. Nucleic Acids Res., 36, (16):e105..

38. Wang, W., Zheng, H.K., Fan, C.Z., Li, J., Shi, J.J., Cai, Z.Q., Zhang, G.J., Liu, D.Y., Zhang, J.G., Vang, S. et al. (2006) High rate of chimeric gene origination by retroposition in plant genomes. The Plant cell, 18, 1791–1802.

39. Zou, C., Lehti-Shiu, M.D., Thibaud-Nissen, F., Prakash, T., Buell, C.R. and Shiu, S.H. (2009) Evolutionary and expression signatures of pseudogenes in *Arabidopsis* and rice. Plant Physiol., 151, 3–15.

40. Matsuba, Y., Nguyen, T.T.H., Wiegert, K., Falara, V., Gonzales-Vigil, E., Leong, B., Schafer, P., Kudrna, D., Wing, R.A., Bolger, A.M. et al. (2013) Evolution of a complex locus for terpene biosynthesis in *Solanum*. The Plant cell, 25, 2022–2036.

41. Mostovoy, Y., Levy-Sakin, M., Lam, J., Lam, E.T., Hastie, A.R., Marks, P., Lee, J., Chu, C., Lin, C., Dzakula, Z. et al. (2016) A hybrid approach for de novo human genome sequence assembly and phasing. Nature Methods, 13, 587–590.

