## Supplementary Figures for "Read coverage as an indicator of misassembly in a short-read based genome assembly"

### SUPPLEMENTARY TEXT

#### Sensitivity and accuracy of CNVnator in RD value calculation

To evaluate the sensitivity and accuracy of CNVnator in RD calculation, we resampled reads from tomato genome based on simulated RDs, where i) the only possible RD values were 0 (LC), 1 (BG), or 2 (HC) (**Fig. S6A,B**); ii) the analysis RDs were discretized to integers (**Fig. S6C**); iii) analysis RDs were used (**Fig. S6D**). The resultant RDs were highly correlated with the known, simulated RDs (PCC≥0.97), suggesting that detection of HC/LC/BG region using CNVnator is reasonable and robust.

#### Impact of genome coverages on RD values

As shown in Fig. 1A, subset of dataset2 at ~30-fold coverage had very similar RD distribution as dataset2. To assess the extent to which read coverage impacts the detection of HC/LC/BG regions, we randomly resampled reads at 5-fold, 10-fold, 20-fold coverages from dataset2. The resultant RDs of all subsets of dataset2 (5~30-fold coverage) were compared to analysis RDs using all reads in dataset2 (~46-fold coverage). The correlation decreased as the read coverage decreased (PCC from 0.99 to 0.89; **Fig. S6E-H**). Genome sequencing reads at ≥20-fold coverage may provide very similar information for HC/LC/BG region detection (PCC=0.98, $\rho \geq 0.92$), while reads at 10- or 5-fold coverage likely have some error rates in HC/LC/BG region detection (PCC ≤ 0.95, $\rho \leq 0.86$). These results also suggest that the RD variation in detected HC/LC/BG regions reflect potential assembly issues or sequencing bias, instead of being noise introduced by random sampling of reads. To test this, a fake dataset, where the reads were randomly sampled from tomato genome to ~46-fold coverage, was used, and as expected, there is no LC or HC region detected.

In addition, to address the impact of artifacts produced by read mapping using BWA-MEM on RD values, the original reads (~46-fold) were re-mapped to the Short-read assembly and the resultant RD values were almost the same as original values (both PCC and $\rho$=1.0; **Fig. S6I**), suggesting that the impact, if any, is negligible.

#### Choice of q-value threshold in HC/LC/BG region detection

To get HC/LC/BG regions with high confidence, regions with q0 (proportion of reads with multiple matches across genome in a region) >=0.5 (1) were filtered out because they likely represented repetitive sequences. F measure (F1) values were used to measure the consistency between true HC/LC/BG region designations and new HC/LC/BG regions determined using resampled reads. F1 were calculated as: F-measure = $\frac{2*precision*recall}{precision+recall}$, where precision = $\frac{TP}{TP+FP}$; recall = $\frac{TP}{TP+FN}$; TP = true positive, FP = false positive, FN = false negative (**Fig. S7**). P-values of identified HC and LC regions were adjusted to account for multiple testing (2). To choose an adjusted p-value (q-value) to maximize F1 scores, HC and LC regions identified using reads resampled by three strategies and reads at 30-fold coverage as **Fig. S6** were compared to HC and LC regions detected using dataset2. F1 score varies when different q-values were used as thresholds to call HC/LC/BG regions (**Fig. S7**). In **Fig. S7A-C** and **E-G**, F1 fitted curve arrives a platform after q-value > 0.06, whereas in **Fig. S7D and H**, the break point of q-value is 0.08. Therefore, only regions with q-value < 0.08 were retained, resulting in 1227 HC and 15095 LC regions.

### REFERENCES


1. Abyzov,A., Urban,A.E., Snyder,M. and Gerstein,M. (2011) CNVnator: an approach to discover, genotype, and characterize typical and atypical CNVs from family and population genome sequencing. *Genome Res.*, **21**, 974–984.
2. Benjamini,Y. and Hochberg,Y. (1995) Controlling the False Discovery Rate: A Practical and Powerful Approach to Multiple Testing. *J. R. Stat. Soc. Series B Stat. Methodol*, **57**, 289–300.


### SUPPLEMENTARY TABLES

**Supplementary Table S1. Choice of bin size**
**Supplementary Table S2. Performance of 3-class or binary prediction models**
**Supplementary Table S3. Importance values and Kruskal test of features in Model 34**
**Supplementary Table S4. Correlation among densities of HC/LC/BG regions and genomic features**
**Supplementary Table S5. Importance values of features in Model 35**
**Supplementary Table S6. Importance values of features in Model 34B**
**Supplementary Table S7. Importance values of features in Model 35B**
**Supplementary Table S8. Number of annotated sequences in HC regions and 10,000 randomly sampled BG regions (based on the number and length distribution of HC regions)**
**Supplementary Table S9. Gene set enrichment analysis for Pfam domains**
**Supplementary Table S10. Gene set enrichment analysis for GO terms**

**Supplementary Table S11. Gene set enrichment analysis for metabolic pathways**

**Supplementary Table S12. Aligned regions between Short-read assembly and Long-read assembly**

**Supplementary Table S13. Categories of regions in Short-read assembly**

**Supplementary Table S14. Corresponding variable coverage regions between two assemblies**

**Supplementary Table S15. HC regions in Short-read assembly and corresponding variable coverage regions in Long-read assembly**

**Supplementary Table S16. Importance values of features in Model 37**

**Supplementary Table S17. Importance values of features in Model 38**

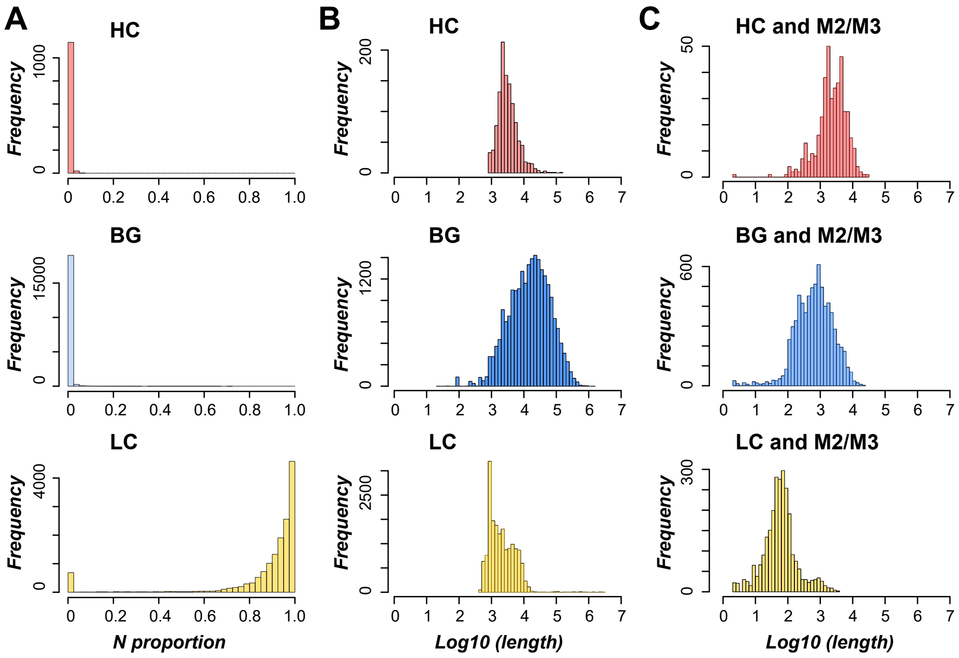


**Figure S1. Properties of HC/LC/BG regions with high confidence.** (**A)** Proportion of Ns in HC/LC/BG regions. (**B**) Length distribution of HC/LC/BG regions. (**C**) Length distribution of overlapped regions between HC/LC/BG and M2/M3 regions.

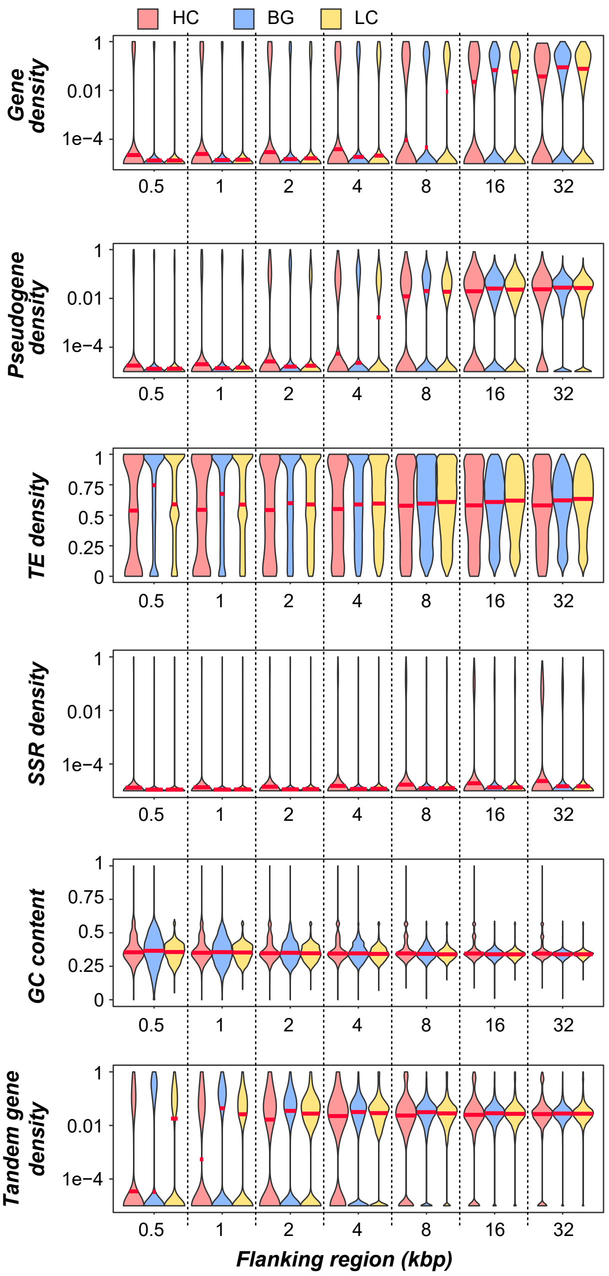


**Figure S2. Genomic feature distributions in flanking regions of HC/LC/BG regions of different length (0.5~32Kb).** Violin plots showing distributions of GC content, and densities of genes, tandemly duplicated genes, pseudogenes, transposable element and SSRs in HC, BG, and LC regions. Red line indicates median value. Statistics analysis was conducted using Wilcoxon rank sum test, *** indicates $p$<1e-3.

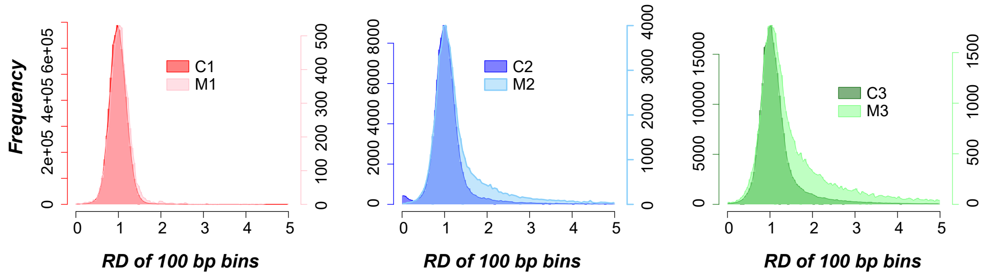


**Figure S3. RD distribution of 100 bp BG bins overlapped with aligned region categories.** For each plot, left Y-axis: number of BG bins in correctly assembled category (C1, C2 or C3); right Y-axis: number of BG bins in mis-assembled category (M1, M2 or M3). The categories are defined in **Fig. 5A**.

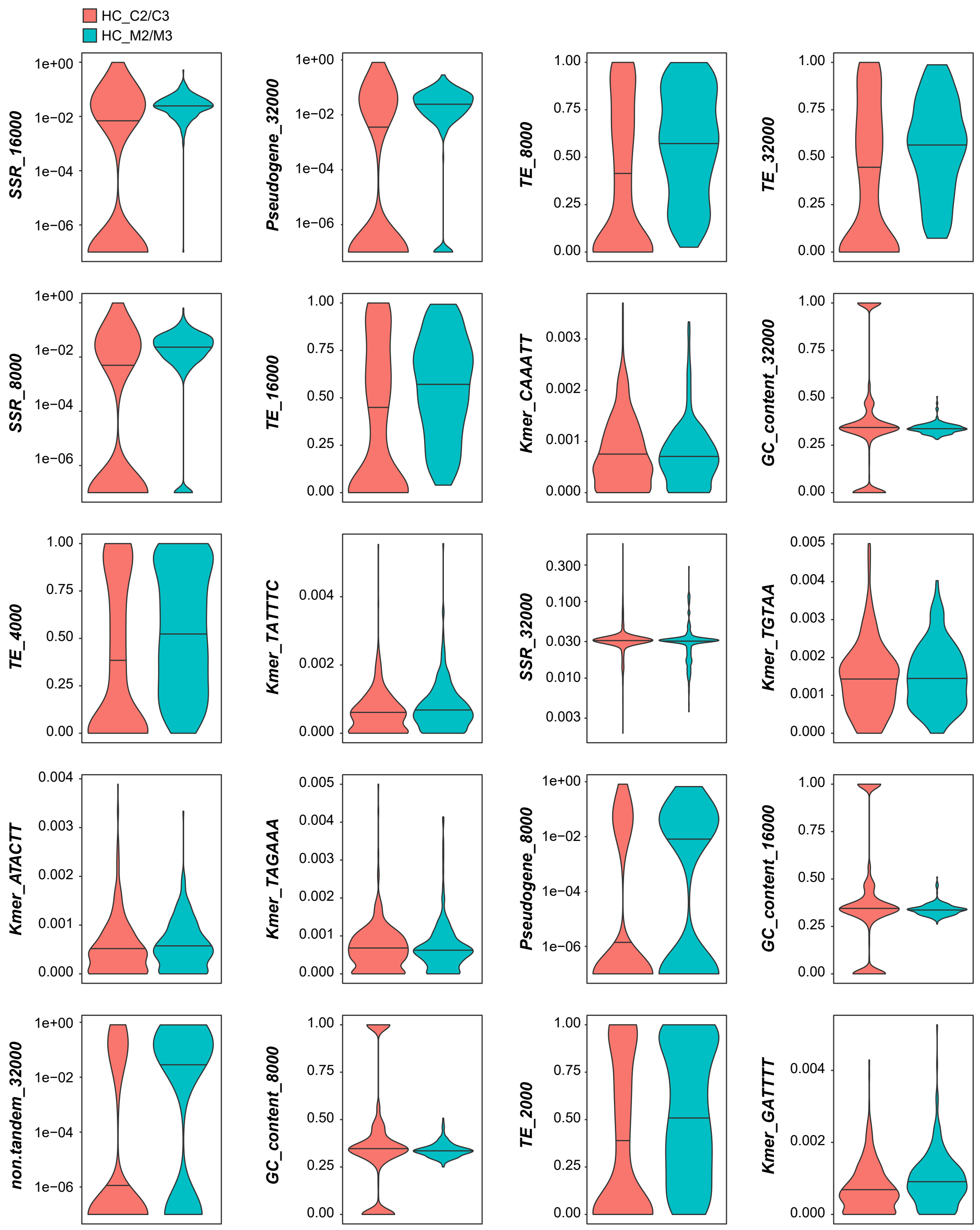


**Figure S4. Important features in model distinguishing HC_M2/M3 and HC_C2/C3.** Violin plots show distributions of each features or GC contents within regions or in flanking regions. Lines within violin plots indicate median values.

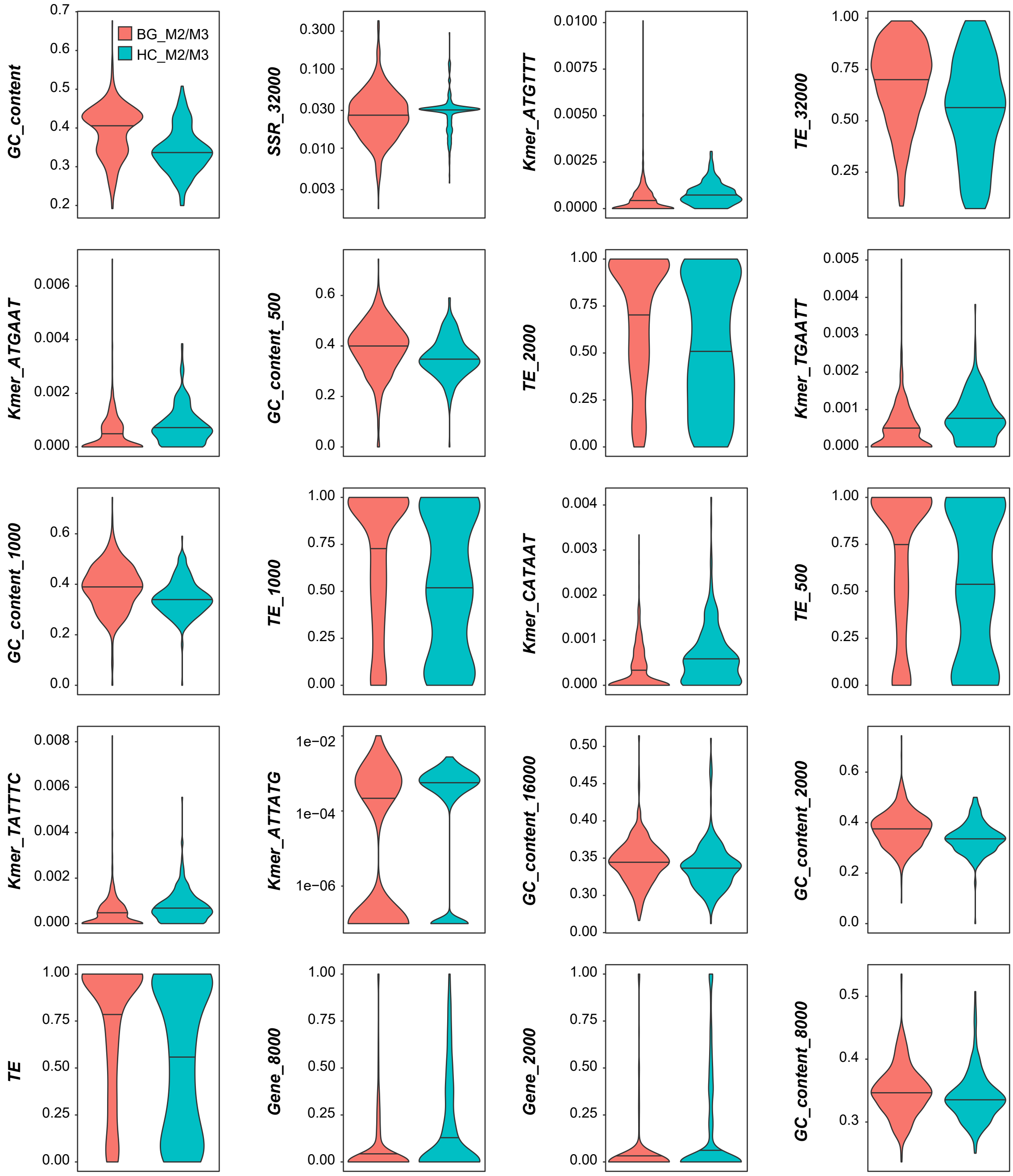


**Figure S5. Distributions of important features in model distinguishing HC_M2/M3 and BG_M2/M3.** Violin plots show distributions of each features or GC contents within regions or in flanking regions. Lines within violin plots indicate median values.

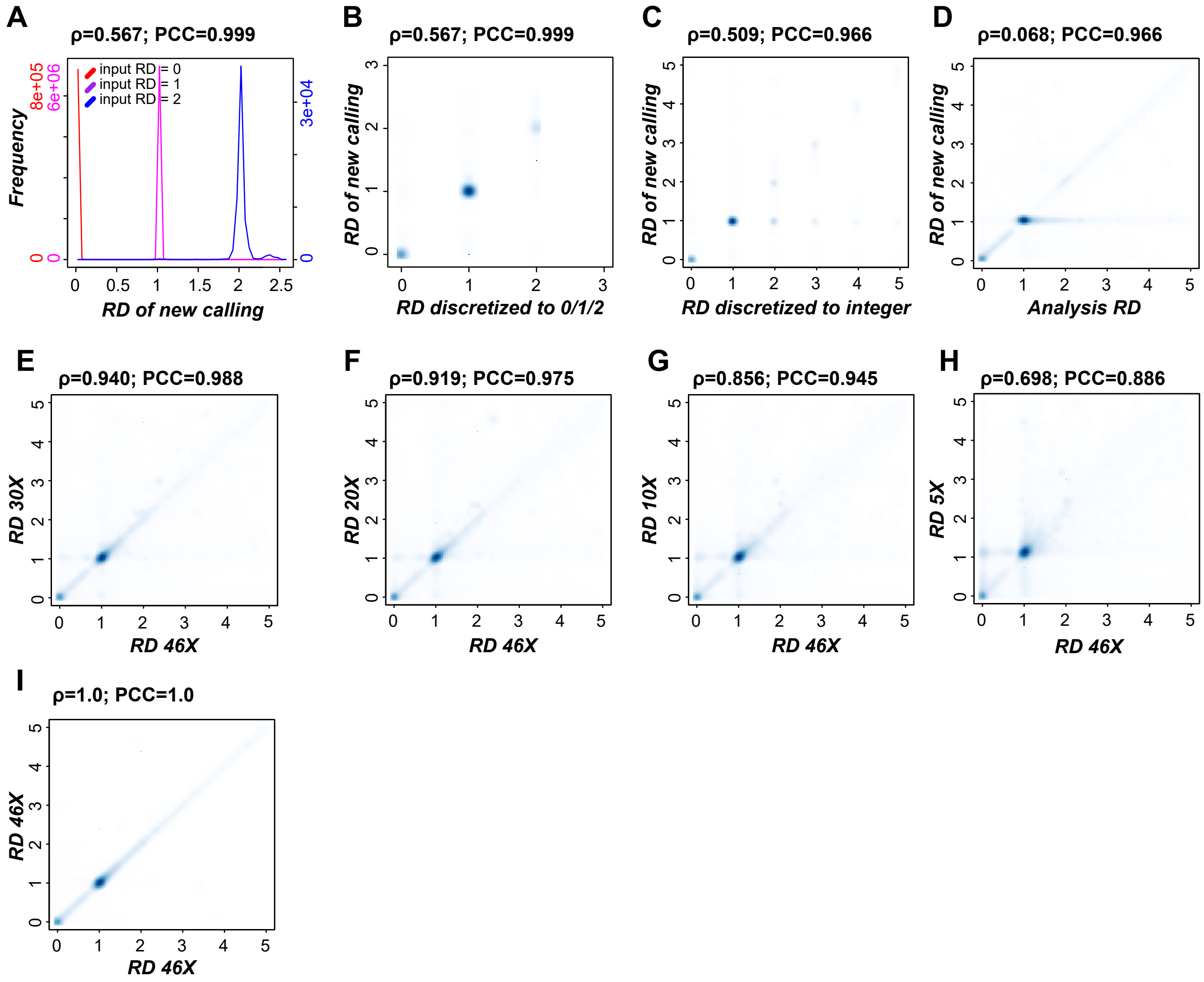


**Figure S6. Sensitivity and accuracy of CNVnator in RD value calculation, and impact of genome coverages on RD values.** (**A**) RD distributions of new CNVnator runs by mapping re-sampled reads from the tomato genome based on simulated input RD, where the only possible RD values were 0 (LC), 1 (BG), or 2 (HC). (**B, C, D**) Correlation between known, simulated input RDs and new RD values from new CNVnator run using the resampled reads. In (**B**), the simulated RD values were generated as in (**A**). In (**C**), the analysis RD values (those generated with CNVator by mapping dataset2 reads on to the tomato genome, see **Methods**) were first discretized (rounded) to their closest integers, then the rounded RD values were used for resampling reads for determining new RD values. In (**D**), the analysis RD values were directly used for resampling reads for determining new RD values. (**E-I**) Correlation between RD values using all reads (46X coverage) and RD values using subsets of reads at variable coverage: (**E**) 30X, (**F**) 20X, (**G**) 10X, (**H**) 5X, (**I**)46X.

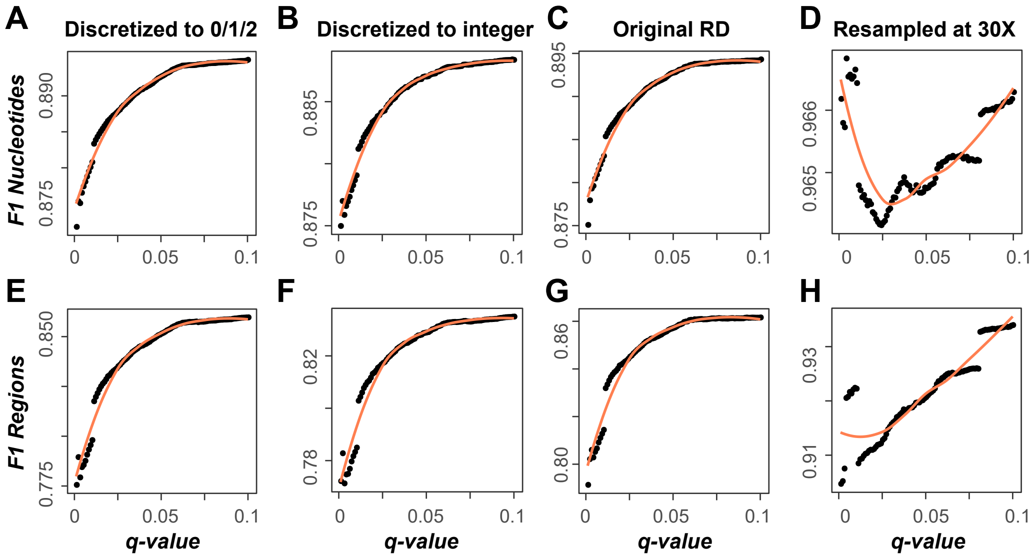


**Figure S7. F1 of HC/LC/BG region calling.** (**A-D**) CNVnator runs were conducted using resampled reads based on different starting RD values as in Figure S3 using read dataset 2. As in Figure S3, the simulated RD values include: (**A**) possible RD values of only 0 (LC), 1 (BG), or 2 (HC); (**B**) analysis RD values discretized/rounded to integers; (**C**) analysis RD without discretization; (**D**) analysis RD without discretization but down-sampled to 30X (note the y-axis range is much smaller compared to (**A**)-(**C**)). Each dot indicates an F1 value (y-axis) at a given q-value threshold (x-axis), where F1 was calculated using: (1) numbers of nucleotides in overlapping regions between the HC/LC region designations based on the analysis RD and new HC/LC regions determined using resampled reads (True Positive), (2) numbers of nucleotides in true HC/LC regions but determined as BG regions in new run (False Negative), and (3) numbers of nucleotides in true BG regions but determined as HC/LC regions in new run (False Positive, see **Supplementary Text**). Orange line: LOESS fitted curve. (**E-H**) Same as (**A-D**) except that the F1 was determined based on numbers of regions as opposed to numbers of nucleotides.
